# misosoup: A metabolic modeling tool for identifying minimal microbial communities provides valuable insights into microbial ecology and biotechnological applications

**DOI:** 10.1101/2025.08.07.669121

**Authors:** Nicolas Ochsner, Magdalena San Román, Adrián Jiménez-Fernández, Sebastian Bonhoeffer, Alberto Pascual-García

## Abstract

Microbial survival and function often depend on metabolic interactions within communities. Therefore, a central question in disentangling microbial organization is determining which minimal groups of species are able to thrive in a given medium – referred to as “minimal communities”. Answering this question is essential for understanding microbial distribution, enhancing laboratory cultivation, and designing synthetic communities (SynComs). Here, we introduce `misosoup`, a Python package for identifying minimal communities (MInimal Supplying community Search). Through genome-scale constraint-based metabolic modeling, `misosoup` enables the systematic identification of communities that support microbial growth in environments where individual species fail to survive alone. We validate `misosoup` against experimentally verified minimal communities, demonstrating its ability to predict known cooperative interactions, cocultures, and consortia with biotechnological potential. We further illustrate the use of `misosoup` to investigate broad microbial ecology questions by applying it to a set of 60 marine microbes, finding pervasive cross-feeding-driven niche expansion, and showing how the detailed outputs provided by `misosoup` facilitate research on hot topics such as the identification of functional groups. In summary, `misosoup` provides a powerful tool for microbial ecology and community design, with potential applications in both research and biotechnological innovation.

**Importance:** Microbes often rely on each other to survive, especially in environments where they can’t live alone. Understanding which small groups of microbes can thrive together—called minimal communities—is key to improving lab research, designing synthetic ecosystems, and exploring how microbes spread in nature. To support this, we developed `misosoup`, a Python tool that identifies these communities using advanced metabolic modeling. `misosoup` helps scientists discover how microbes cooperate by sharing nutrients, a process known as metabolic cross-feeding. When tested on sets of species from different origins, the tool showed that species could thrive in more environments when part of a group. This highlights the importance of teamwork in microbial life. `misosoup` not only predicts these interactions but also provides detailed insights that can guide ecological studies and biotechnological innovation. By revealing how microbes support each other, `misosoup` contributes to a deeper understanding of life’s interconnectedness and offers tools for solving real-world challenges.

## 1 Introduction

Microorganisms in nature form highly diverse communities shaped by intricate biotic interactions. While community members often compete for space and resources, they also depend on one another for essential metabolites, such as signaling molecules, siderophores, amino acids, vitamins, and for mitigating the effects of toxic compounds [1]. Microbial communities also play pivotal roles in biotechnological applications, including bioremediation, biofuel production, and pollutant biotransformation. Identifying the necessary and sufficient partners that enable species to coexist within these communities is essential for both ecological insight and the development of simplified synthetic communities.

Simplified synthetic communities (generically termed consortia or SynComs) open new opportunities to handle experimentally-tractable systems for the study of microbial interactions and the design of microbial communities delivering specific services. Different strategies have been proposed to build these communities, being the ‘bottom-up’ approach –in which isolated strains are combined to build the consortia– the most prevalent [2, 3, 4]. The ‘bottom-up’ approach, however, faces some challenges. It requires the prior isolation of each species, restricting the experiment to culturable organisms only [5]. In addition, the number of possible community combinations increases exponentially with the number of species, making it impractical to culture all potential communities. Therefore, a critical challenge in constructing novel consortia lies in identifying stable combinations of strains that can coexist within a given medium. While recent advancements, such as the development of innovative co-culture plates [6], microfluidic chips [7], and cost-effective protocols for assembling combinatorially complete communities [8], have significantly expanded experimental capabilities, these approaches still face the inherent limitations of traditional “bottom-up” methodologies.

A powerful alternative approach to overcome these limitations comes from computational modeling. In particular, since a significant fraction of microbial interactions are of metabolic nature, metabolic modeling stands out as a valuable complementary approach for the study of microbial communities. This computational framework leverages genome-scale metabolic models of species, that encompass all known metabolic reactions within an organism. Further, constraint-based methods, such as Flux Balance Analysis (FBA) [9], enable the determination of growth rates and consumption/secretion patterns of species at steady-state within defined environments. This framework has proven effective in various applications involving clonal populations, including growth prediction, by-product secretion [10], the effect of gene deletions [11], or adaptive evolution [12].

In the context of community-level metabolic modeling, a focus on the search and design of consortia performing specific functions is prevalent, reviewed in [13, 14, 15]. An example of this approach is `FLYCOP` [16] a tool that integrates `COMETS` [17] to explore multiple community configurations, in order to find the community that optimizes a specified goal. Another example is `MinMicrobiome` [18], which uses a top-down approach to identify minimal microbial communities capable of performing a specific function, particularly suited for optimizing short-chain fatty acid secretion from AGORA metabolic models. Given the limitations of constrained-based metabolic modeling to handle a large number of strains, graph-theoretic approaches are proposed. For example, `CoMiDA` [19] finds the smallest set of microbial species that collectively possess the enzymatic capacity to synthesize a desired target product, a strategy later followed in `Metage2Metabo` [20].

In addition, metabolic modeling has proven to be a powerful tool in studying microbial ecology. Both constrained-based and graph-theoretic models have been employed to explore ecological interactions [21, 22, 23]. An important step towards community-level modeling was the work of Klitgord and Segré who developed an algorithm to find the minimal medium in which pairs of strains could coexist [24]. This work inspired other constrained-based methods that extended this search to larger communities [25], and later to evaluate the potential for cooperation and competition of specific consortia implemented in SMETANA [26]. These tools facilitate the exploration of broad questions such as the relative importance of cooperation and competition in terrestrial [27] and marine systems [28], or the development of an atlas of putative costless secretions [29].

In this work, we contribute towards the development of community-level metabolic modeling by focusing on the problem of species coexistence. We ask which minimal communities can coexist given a set of species and a predefined medium. A minimal community is one in which every species is necessary and sufficient for the survival of all members in the community. We differentiate between *complete minimal supplying communities* (CMSCs), in which no subset of species in the community can thrive in that medium, and *minimal supplying communities* (MSCs), which are solutions associated with a particular focal strain. Here, the focal strain is defined as a strain that cannot grow on the selected medium in isolation and therefore requires metabolite supply from partner strains to succeed. In a MSC, although the focal strain depends on its suppliers for growth, subsets of the supplying partner strains may still be able to grow independently in the medium. Therefore, if each member of a CMSC is considered in turn as the focal strain, the CMSC itself must appear as one of the corresponding MSC solutions-hence the term ‘complete’.

To address this question, we built on top of methods established in SMETANA to develop an algorithm for the MInimal Supplying cOmmunity Search (`misosoup`), implemented as a Python package. A schematic representation of the problem addressed, and its implementation is presented in Fig. 1. Unlike other methods focusing on finding a minimal community with a prespecified function (e.g. `MinMicrobiome`) or the potential metabolic exchanges of a specific community (`SMETANA`), `misosoup` solves a mixed-integer linear programming (MILP) problem to enumerate all feasible minimal communities for a given set of strains and a specific medium.

**Figure 1:**
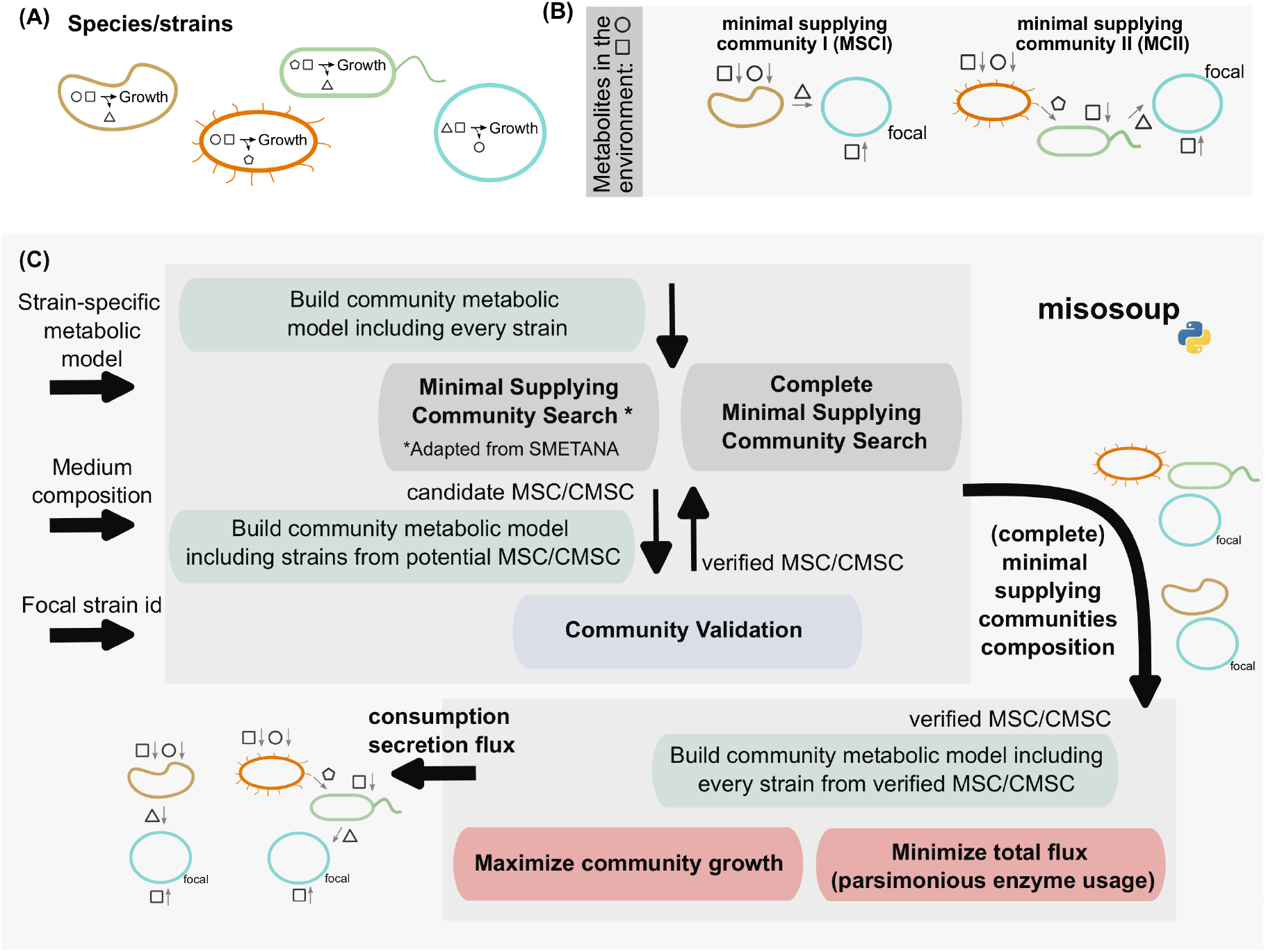
Minimal supplying communities and misosoup. (A) Representation of hypothetical bacterial strains with shapes representing metabolic requirements and secreted metabolites. (B) Alternative MSCs (I and II) for these strains in an environment with two resources (square and circle), where the blue strain is focal. MSCI and MSCII are minimal as removal of any species leads to the focal species’ extinction. (C) Explanation of `misosoup`, its inputs and outputs. The identification process involves two consecutive optimizations: the first step solves a mixed-integer linear programming (MILP) problem to find candidate MSCs, and the second step uses linear programming (LP) to verify that the identified communities satisfy all imposed constraints. Once community feasibility is tested, different optimizations are available to find the consumption and secretion fluxes of every community member.

Crucially, `misosoup` adds an evaluation step using linear programming (LP) where every minimal community identified is tested to exclude solutions affected by numerical instability, inherent to complex optimizations. In addition, `misosoup` also offers optional optimizations, such as maximizing community biomass or minimizing total flux, which provide detailed consumption and secretion rates for each strain in the community.

We illustrate the capacity of `misosoup` by considering different examples and quality of metabolic models. Firstly, we tested `misosoup` on a set of experimentally-verified minimal communities with curated metabolic models, finding that it accurately predicts the strains coexisting under the conditions defined experimentally [21]. Next, we considered a set of custom metabolic models to show that `misosoup` can find MSCs of biotechnological interest.

To continue, we studied the ability of `misosoup` to provide insights into the capacity of predicting growth of auxotroph strains in coculture experiments [30], with semi-curated AGORA models, a strategy that could be used to grow isolates in media in which they can’t grow. We finally move on to microbial ecology, considering a larger set of strains and draft metabolic models to explore broader questions, such as the prevalence of cross-feeding-driven niche expansion or the possibility of finding functional groups as building blocks of microbial biodiversity [31, 32].

In summary, `misosoup` is a versatile tool that can be used to aid in organism cultivation, the design of microbial communities, and we demonstrate its applicability to microbial ecology. Software and documentation of `misosoup` are freely available at https://sirno.github.io/misosoup.

## 2 Results

### 2.1 misosoup correctly predicts experimentally-verified minimal communities

`misosoup` is designed for large-scale community searches involving many strains and/or media conditions. However, experimentally verified consortia for which curated metabolic models are available remain scarce in the literature, limiting the availability of suitable ground truth datasets. For this reason, and as a proof of concept, we first evaluated a small set of experimentally validated two- and three-strain communities. Although this is far below the scale for which Misosoup is intended, these consortia provide reliable validation cases.

The first community (Fig. 2A) consisted of two mutant *Escherichia coli* strains auxotrophic for leucine (*E. coli* Δ*leuA*) and lysine (*E. coli* Δ*lysA*) respectively. In glucose minimal medium, the strains grow only when they exchange the essential metabolites leucine and lysine [33]. `misosoup` was able to predict the coexistence of both strains when they were grown in coculture, and their inability to grow in isolation. Moreover, the exchanges found were consistent with the expectation, with *E. coli* Δ*leuA* secreting lysine that is consumed by *E. coli* Δ*lysA*, and the other way around for leucine.

**Figure 2:**
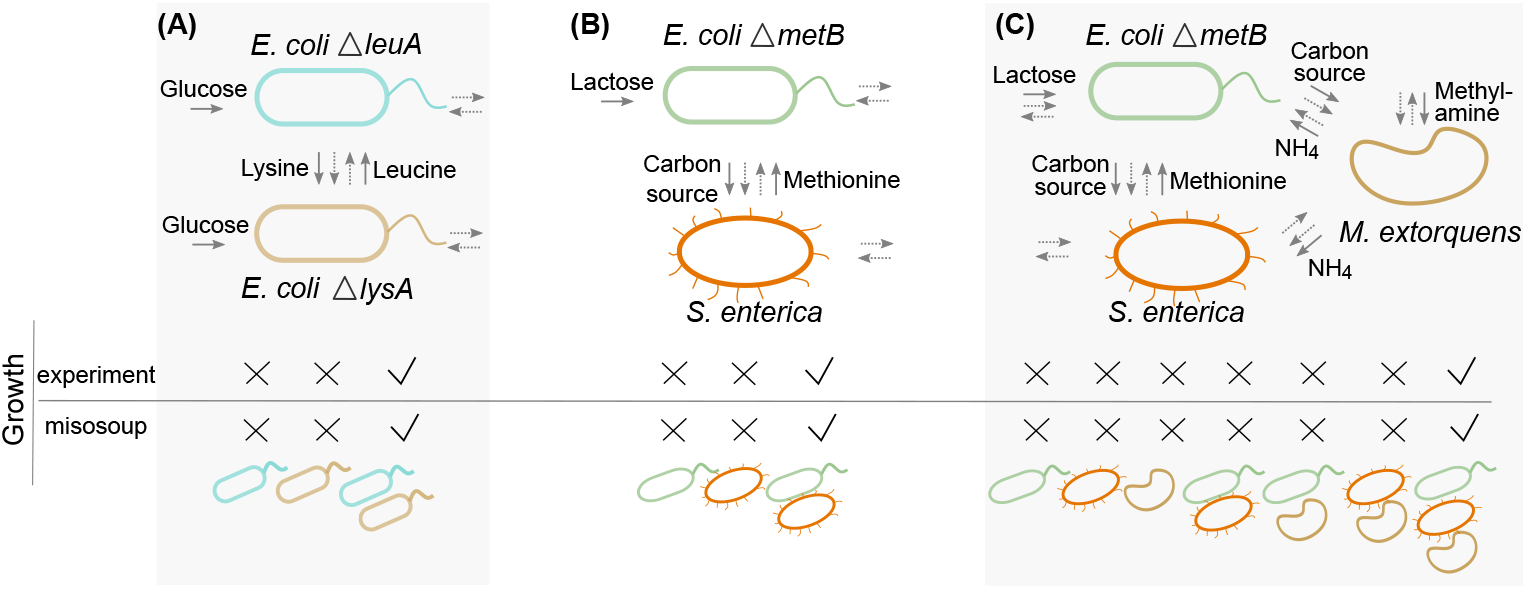
(A) Computationally predicted and experimentally tested minimal communities. The strains’ consumption/secretion pattern, as predicted with `misosoup`, is indicated with arrows: main consumed/secreted metabolites are indicated with arrows and the metabolite name; dashed arrows indicate that `misosoup` also predicts the consumption/secretion of other metabolites. `misosoup` correctly predicts that two (A and B) or three (C) strains are required for growth in glucose minimal medium (A), lactose minimal medium (B) and lactose minimal medium with methylamine as nitrogen source (C).

The second community (Fig. 2B) consisted of a methionine auxotroph mutant *E. coli ΔmetB*, and *Salmonella enterica* mutant strain able to secrete methionine. *S. enterica* cannot grow in lactose, hence relying on the *E. coli* strain [21]. `misosoup` correctly predicted coexistence in lactose minimal medium when ammonium was the nitrogen source, further confirming that both strains were unable to grow independently. The exchange of alternative C-sources that *S. enterica* consumed and of methionine, required by *E. coli*, were also correctly predicted.

In addition, we verified that, when ammonium was substituted by methylamine as nitrogen source, both strains were unable to grow in co-culture, consistent with experiments [21]. Following experimental results we asked if it is therefore possible to create a 3-species obligatory mutualistic consortium by introducing a species able to assimilate methylamine as a nitrogen source. *Methylobacterium extorquens* is able to use methylamine as both nitrogen and carbon source. A mutant lacking hydroxypyruvate reductase (Δ*hprA*) cannot assimilate carbon from methylamine, relying on the acetate excreted by *E. coli* for growth, and can provide nitrogen to the other two species.

`misosoup` correctly predicted that no pairwise combination of these three species had positive growth on lactose minimal medium with methylamine as a nitrogen source, while coexistence was only observed when the three species were grown together (Fig. 2C). Notably, it also correctly predicted the experimentally-observed exchanges [21] as detailed in Fig. 2C.

### 2.2 misosoup finds minimal communities with biotechnological potential

As a first application of `misosoup` we considered finding consortia with biotechnological interest, since an important bottleneck in often finding feasible consortia candidates, a question for which `misosoup` is especially well-suited.

To illustrate this application, we searched for consortia able to produce biogas, considering 9 strains previously used in the publication of the metabolic modelling tool RedCom [35], whose custom metabolic models were available from the publication. Two of the strains were methanogenic archaea (*Methanospir-illum hungatei*, MH; and *Methanosarcina barkeri*, MB), and 6 primary and secondary fermenters (Escherichia coli, EC; *Clostridium acetobutylicum*, CA; *Propionibacterium freudenreichii*, PF; *Acetobacterium woodii*, AW; *Desulfovibrio vulgaris*, DV; *Syntrophomonas wolfei*, SW; and *Syntrophobacter fumaroxidans*, SF) that may generate different compounds from an initial resource such as glucose or ethanol (e.g. acetate, formate) for downstream production of CH4 by the archaea.

To reproduce results found in [35], we focused on minimal communities growing on ethanol as carbon source. We first searched for CMSCs in which no species can grow in isolation in the specified medium, and whose detection remains elusive for other tools. We found two CMSCs (Fig. 3, middle column), one was a cross-feeding consortium formed by DV and MB, and the other one was AW growing in isolation. Note that, although species growing in isolation do not represent consortia, `misosoup` returns them since formally they fulfill CMSC definition, which allows the user to identify what strains can grow in isolation. We termed suchsolutions CMSC singletons.

**Figure 3:**
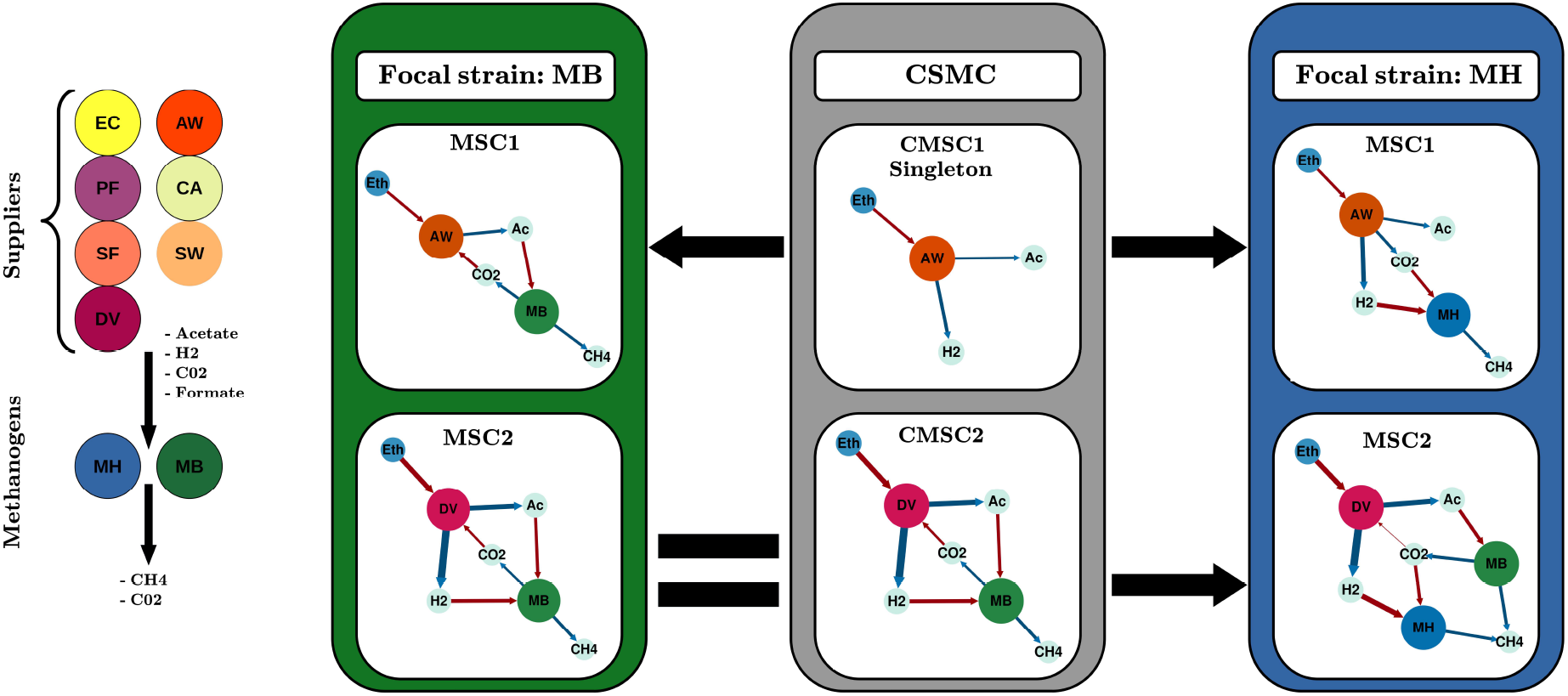
Prediction of biogas-producing consortia. We considered nine strains to identify consortia capable of producing *CH*_4_ from ethanol. Our analysis yielded two candidate minimal consortia (CMSCs; middle column). One of these consisted of AW alone (a “singleton”). When MB was used as the focal strain, we identified one new consortium and one previously reported CMSC (left column). When MH was used as the focal strain, we found two additional consortia (right column). One of these matches a three-species consortium described in the literature.

Since, to produce CH4, at least one of the archeas is needed, next we searched for MSCs, considering each of the archaea as a focal strain. We found that MB was able to form a cross-feeding consortium with AW and, consistently, the CMSC previously found (Fig. 3, left column). When MH was the focal strain, it also formed a consortium with AW but there was no supply from MH to AW, as it was the case in the MB-AW consortia, reflecting a commensal structure (Fig. 3, right column). Notably, we found that DV and MB, the second CMSC, also provided the resources needed for MH to thrive, forming a three-species consortium.

Interestingly, the three-species consortium found was the one reported in RedCom [35]. But, while the authors proposed this consortia because a similar one was previously reported in Ref. [36], `misosoup` effortlessly found it as a solution. Importantly, the analysis also provided insights into how the consortium is organized into subunits, with DV and MB forming the core as a CMSC and MH depending on them.

### 2.3 misosoup predicts large-scale co-culture experiments

The next example considered predicting coculture experiments for species that cannot grow in certain media in isolation. We considered an experiment by Oña *et al* . [30] in which they knocked out a set of strains to generate amino-acids auxotrophs, to then verify their ability to grow in coculture, a situation they termed ‘niche expansion’. Therefore, niche expansion here is a situation wherein an individual organism lacks the capacity for independent growth in a specific abiotic environment but can thrive in that same environment as part of a microbial community.

We selected three strains assayed in the publication (*Bacillus subtilis* 3610, *Escherichia coli* B225113, *Pseudomonas fluorescens* SBW25) that were available in AGORA2, a database of automatically-curated metabolic models [34]. For each strain, we generated four auxotrophic metabolic models, each lacking the ability to synthesize one amino acid (leucine, arginine, tryptophan and histidine corresponding to L, R, W, and H in Fig. 3; see Methods). We then verified their growth in monoculture across 31 experimental media (see Methods), each supplemented with the amino acid that the corresponding auxotroph cannot synthesize. Finally, we gapfilled the models whenever growth was observed in supplemented monoculture experiments but not in the models (see Methods and Suppl. Materials). We should note that `misosoup` ability to predict cocultures depends on the experimental criteria considered to determine that growth was observed (e.g. minimum OD or number of replicates in which growth should be observed) because, in turn, these choices determine model gap-filling. In Suppl. Materials we evaluated the influence of these choices. In the following, we quantified `misosoup`’s ability to predict cocultures restricting the analysis to combinations of media and strains in which individual models of both strains were, after gap-filling, in agreement with the experiments in monoculture.

In Fig. 4 we show `misosoup` coculture growth predictions for pairs of auxotrophs grown on different C sources, with blue cells representing agreement between prediction and experiment. Notably, accuracy was above 0.7 in more than half of the experiments performed, highlighting `misosoup` predictive performance.

**Figure 4:**
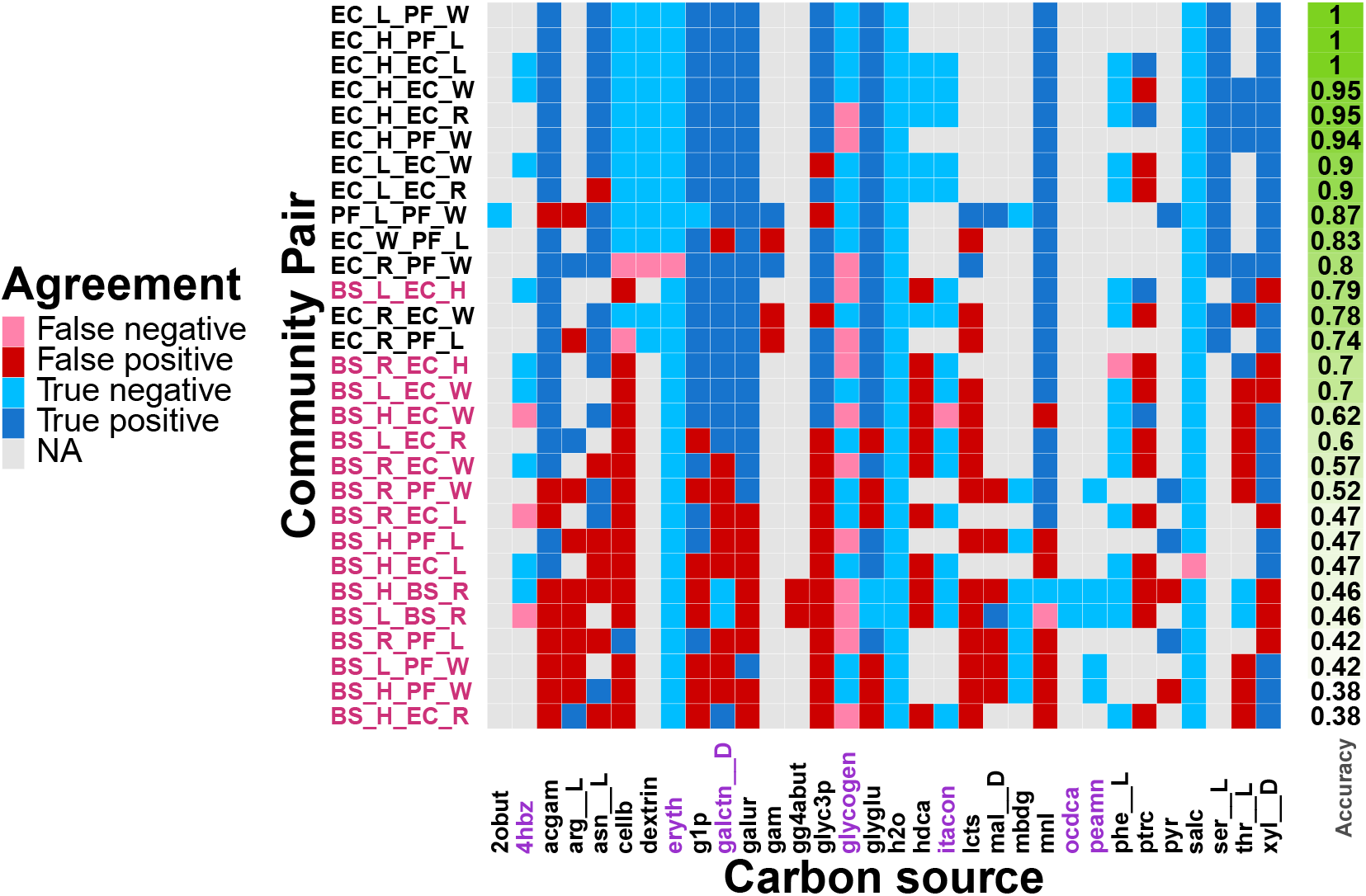
Prediction of coculture experiments. Heatmap representing pairs of auxotrophs cocultured (rows) in different media (columns) with the compound indicated as sole C source. Agreement between `misosoup` and experiments are shown in blue (true positives and negatives), and disagreements in pink (false positives) and red (false negatives). Gray cells represent experiments in which any of the two members had no agreement with the experiments when grown in monoculture, and were excluded from the computation of accuracy (green bar). Labels of co-cultures including a lower quality model (*B. subtilis*) and media that could not be gapfilled are highlighted. These cases explain most discrepancies with model predictions. More details in Main Text.

Importantly, most simulations that did not match the experiment (red/pink cells) are those cocultures in which *B. subtilis* was present (Fig. 4). In Suppl. Materials we show that this model had a poor performance compared to the other two strains, even after gapfilling. Since it tends to generate false positives, it suggests that the model was likely over-gap-filled (see strong increase in sensitivity after gap-filling in Suppl. Fig. S2). Another source of discrepancy can be attributed to media that could not be gap-filled, which either do not show a good agreement in monoculture (e.g. ocdca), or tend to generate false negatives (glycogen).

Finally, `misosoup` also predicted 25 more cocultures not assayed experimentally, one of them involving three strains (Suppl. Fig. S5, and even included auxotrophs from the same species (*Pseudomonas fluorescens*) which were not predicted for the other species. We should, however, be cautious with pairs involving *B. subtilis*, since we observed they were often associated to false positives. Our results highlight the potential of `misosoup` to find conditions in which strains can grow, opening new avenues for culturomics experiments.

### 2.4 misosoup application in a microbial ecology experiment supports pervasive cross-feeding-driven niche expansion

Next, we illustrate the potential of `misosoup` to explore broad ecological questions. The strains used in this study were isolated from previous experiments in which natural marine communities assembled on synthetic beads uniformly covered with one or two naturally-occurring polymers, mimicking marine snow [37, 38] . reproducible ecological succession was observed to occur on these particles, and it was possible to classify strains as early colonizers, late colonizers or generalists depending on the stage in which they were observed throughout the succession [38]. Interestingly, it was suggested that the capacity of late colonizers to thrive on particles depends on the byproducts produced by early colonizers.

We asked if the succession can be rationalized as an example of niche expansion by investigating a set of 60 marine bacterial strains isolated from these experiments ([37, 38]), searching for MSCs assayed in 22 different environments. We defined the *fundamental niche* as the number of environments in which a strain can grow in isolation and the *realized niche* as the number of environments in which a strain can grow in isolation and in communities. The difference between the realized and fundamental niche constitutes the *niche expansion* (see Methods).

We reconstructed genome-scale draft metabolic models for these strains, using the package CarveMe [39], and simulated growth of each strain in isolation with FBA and in communities with `misosoup` in 22 different environments (see Methods). The environments simulated marine broth and differed only in the carbon source available. We selected carbon sources to be carbohydrates, amino-acids or carboxylic acids, in similar proportions (Fig. 5). See Methods and Suppl. Materials for a detailed description of the media composition.

**Figure 5:**
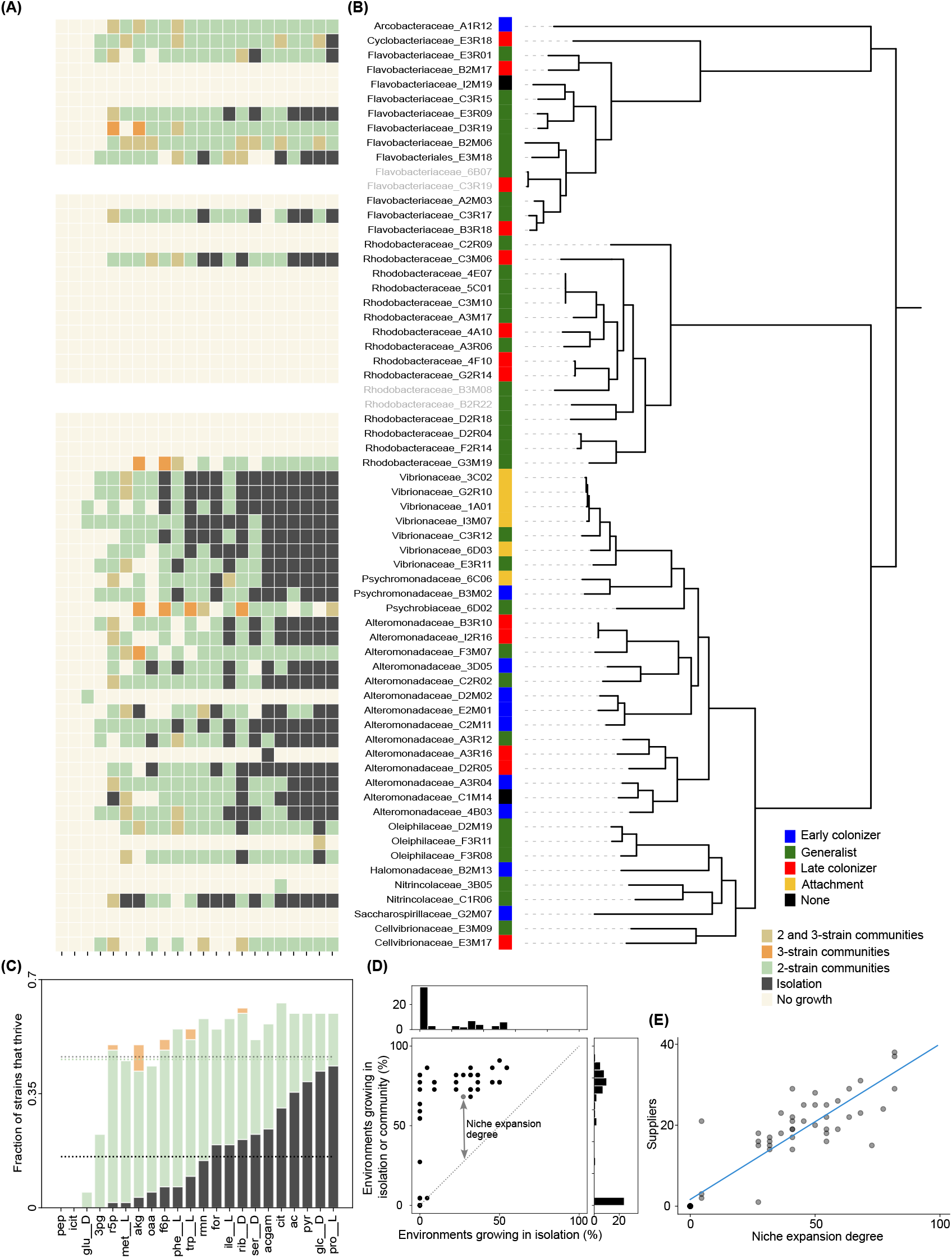
(A) Heatmap illustrating whether a strain thrives in a given carbon source when grown in isolation or in communities consisting of 2 or 3 strains. Strain names correspond to those shown in the tree in panel (B), and carbon sources correspond to those in the graph in panel (C). (B) Phylogenetic tree and ecological strategies of the strains, as inferred in [38]. (C) Fraction of strains thriving in each of the 22 tested environments—either in isolation (black bars) or in communities of 2 (green bars) or 3 (orange bars) strains. The x-axis indicates the short name of the carbon source added to the simulated marine medium (see Suppl. Materials for the full names of the metabolites). Dashed lines represent the average fraction of strains thriving in isolation (black), in communities of 2 strains (green), or in any community size (gray). (D) Relationship between the percentage of environments colonized in isolation or communities and those colonized in isolation only. Each dot represents one of the 60 strains. The arrow highlights the degree of niche expansion for the strain shown in gray. Niche expansion is quantified as the difference between environments colonized in communities or in isolation and those colonized in isolation. (E) Number of different partners each strain has across all solutions facilitating in which niche expansion was observed, and the degree of its niche expansion.

When grown in isolation, we observed significant variation in the fundamental niche sizes of the strains, with closely related strains exhibiting more similar fundamental niches (Fig. 5A and B, and Suppl. Fig. S6). On average, strains were capable of growing in 3.5 environments. Moreover, 32 strains had a fundamental niche size of zero, indicating that they were unable to colonize any of the tested environments (Fig. 5A) with 42% of these strains belonged to the Rhodobacteraceae family (Fig. 5B). Notably, we found that fundamental niche size was positively correlated with both the number of genes (Pearson s *r*^2^ = 0.45) and reactions (*r*^2^ = 0.57 Suppl. Fig. S7) included in the species metabolic models.

Next, we used `misosoup` to search for MSCs selecting each strain at a time as focal strain, while the other 59 strains could make up the supplying community. `misosoup` found a total of 2210 MSCs. 13% of which were made up of three strains while the remainder were made up of only two strains. The analysis of these communities shows that the fraction of strains that can colonize an environment increases if we allow strains to assemble into communities. This is true for all environments tested other than those with `pep` and `icit` in which no strains grew in isolation nor in communities.

On average, 10±9 strains (16±14%) grow in an environment in isolation (black line in Fig. 5C), while 29±12 strains (47±19%) grow either in isolation or as part of a community (gray line in Fig. 5C). As a consequence, 41 strains (66%) show some degree of niche expansion, with an average niche expansion degree of 31% (Fig. 5D). This means that strains can colonize, on average, 7 more environments in communities than in isolation and even those strains that colonize a large fraction of niches in isolation exhibit some degree of niche expansion.

Importantly, we found a significant linear relationship between the degree of niche expansion and the number of different partners required for such expansion (Fig. 5E), which means that the more they expand, the more partners they need. Interestingly, the average phylogenetic relatedness of members within the same MSC is above the family level but below the relatedness expected by random pairing (see Suppl. Fig. S8A), and they also have a significant complementarity in the reactions they host with respect to random pairs (see Suppl. Fig. S8B and C).

Our analysis identified significant variation in how frequently different strains acted as suppliers within these communities. Consistent with previous observations, the most recurrent suppliers were members of the *Vibrionaceae* and *Alteromonadaceae* families, predominantly associated with the attachment and early colonization stages, respectively (Suppl. Fig. S9). Notably, these strains primarily supplied resources to members of the *Flavobacteriaceae* family, classified as generalists.

Overall, we see that metabolic interactions between community members make most niches accessible to a larger number of strains. Further, this enlarged accessibility is aligned with the ecological strategies identified from experiments dominating early stages of the ecological succession and facilitating the later stages.

### 2.5 misosoup helps to explore the architecture of microbial biodiversity

Extensive work is needed to elucidate the mechanisms underpinning niche expansion in experimental settings akin to those conducted in previous studies [30, 40]. The outputs of `misosoup` provide insights into the consumption and secretion of individual community members, which may allow us to elucidate the mechanisms underlying niche expansion, and its consequences in biodiversity maintenance.

Given a repertoire of exchanged metabolites, various consumption and secretion rates (referred to hereafter as flux distributions) may still support species growth, posing computational challenges for exploring every alternative leading to niche expansion. Given these complexities, here we concentrate on a single community flux distribution that yielded maximal community growth-the summation of species’ individual growth rates (see Methods and Suppl. Materials for insights into how other flux distributions could be explored using the output of `misosoup`.)

We investigated the mechanisms giving rise to niche expansion in the focal species of MSCs. To achieve this, we examined how focal strains and their supplier partners utilized the available carbon source. To this end, we categorized each community flux distribution into six distinct classes (Fig. 6A): (i, ii) the focal strain consumes the carbon source; (iii, iv) a supplying strain consumes the carbon source; or (v, vi) both strains share the carbon source. Additionally, we assessed whether the focal strain secreted metabolites subsequently consumed by supplying strains (ii, iv, vi) or not (i, iii, v).

**Figure 6:**
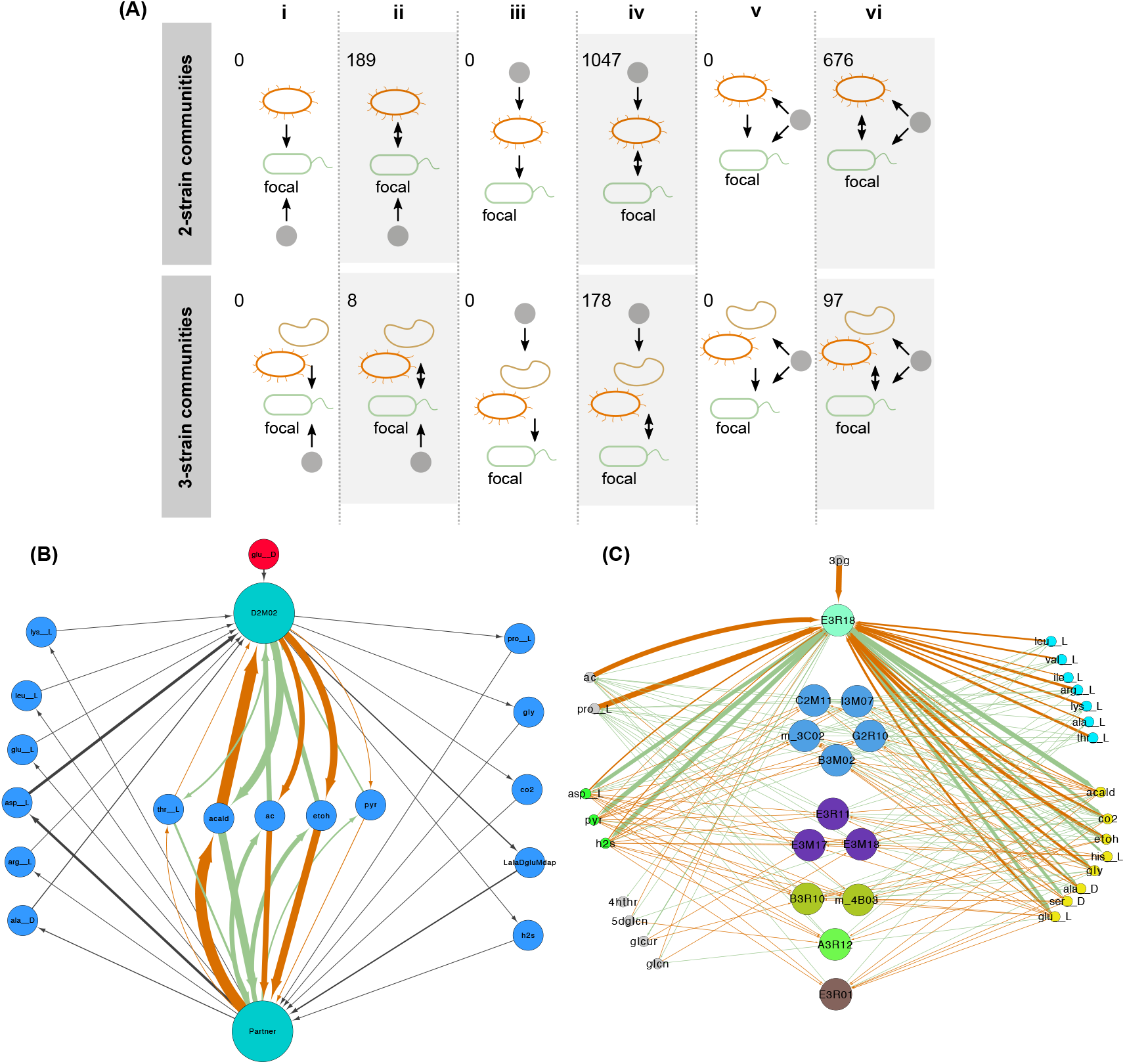
(A) Mechanism giving rise to focal species’ niche expansion in the MSCs. The diagram shows whether communities were made up of 2 or 3 species, and whether they belonged to one of six categories: (i) unidirectional cross-feeding where the focal species consumes the carbon source; (ii) bidirectional cross-feeding where the focal species consumes the carbon source; (iii) unidirectional cross-feeding where a supplying species consumes the carbon source; (iv) bidirectional cross-feeding where a supplying species consumes the carbon source; (v) unidirectional cross-feeding where both species share the carbon source or; (vi) bidirectional cross-feeding where both species hare the carbon source. Numbers on the top left of each panel indicate the number of communities found on each category. (B) The two CMSCs found in `glu_ _D` with common links in black and links specific to each solution highlighted in orange and green. Width is pro portional to the predicted flux with those in black selected randomly from either solution. (C) The 12 CMSCs solutions found in 3pg aggregated. Strains and metabolites are located close in space and with the same color if they belong to the same FG (See Main Text for details). Orange (green) links represent consumption (secretion).

In all MSCs, we observed mutual cross-feeding interactions (ii, iv, and vi in Fig. 6A), wherein the focal strain consumed metabolites secreted by supplying strains and reciprocally secreted metabolites utilized by those supplying strains. Notably, in the majority of communities (56%), the carbon source was consumed by one of the supplying strains, while in 35% of communities, both the focal and supplying strains shared the carbon source. Note that, since the majority of these MSCs were not complete, the supplier1s consumption of metabolites excreted by the focal strain was not a requirement. In the remaining 9% of communities, only the focal strain utilized the carbon source. We did not detect significant differences in these patterns between supplying communities comprising two or three strains.

Next, to illustrate the potential of `misosoup` to explore the architecture of these communities, we investigated two sets of CMSCs found on D-glutamate (`glu_ _D`) and 3-phospho-D-glycerate (`3pg`) with two and 12 CMSCs respectively, all of which were composed of two strains. Exchanged metabolites should be interpreted with caution in these examples since, contrary to other examples presented in previous sections, the presented solutions were obtained with draft metabolic models, and because the problem of finding feasible solutions is highly degenerated. As it was noted above, more biologically and biotechnologically relevant results often rely on setting realistic flux boundaries.

For the two CMSCs identified on `glu_ _D`, strain Alteromonadaceae _D2M02 was a common member in both solutions. This is due to Alteromonadaceae_D2M02 being the only strain capable of metabolizing `glu_ _D`, even though it cannot grow independently and requires a partner. In Fig. 6B we show the union of both CMSCs with the two partners superimposed in a single node. Both solutions exhibit a shared set of metabolites that are consistently consumed and secreted (indicated by black edges). However, a notable observation is a subset of metabolites exhibiting ‘orthogonal’ secretion and consumption patterns: a metabolite consumed by one partner in one solution (orange edges) is precisely what is secreted by the other partner in the alternative solution (green edges). From Alteromonadaceae _D2M02’s perspective, this implies that depending on the specific partner, it can either secrete or consume this particular subset of metabolites, while its interactions with all other metabolites remain consistent across partners.

We also represent the union of 12 CMSCs found on `3pg` (Fig. 6C), where Cyclobacteriaceae_E3R18 (top) appears in all solutions. In this case, Cyclobacteriaceae_E3R18 and another strain (Cellvibrionaceae_E3M09) are the only two strains that can take up `3pg`. We classified strains and metabolites into functional groups using `functionink` [41]. In short, the method classifies strains as belonging to the same functional group if they consume and secrete similar metabolites in their respective solutions (see Methods).

The analysis revealed 5 different functional groups, all of them supplying acetate and proline, that were consumed by Cyclobacteriaceae_E3R18, except for Flavobacteriaceae_E3R01 which only secreted proline (central nodes in Fig. 6C). Interestingly, each group has a distinctive set of amino-acids secreted (cyan metabolites, top right) that were consumed by Cyclobacteriaceae_E3R18. Following groups in the network from top to bottom the amino-acids were Leu, Val, Arg, Thr (blue), Leu, Arg, Lys (violet), Ala_L (light green), Lys, Ala (olive), Lys (maroon). In turn, they uptake different compounds secreted by Cyclobacteriaceae_E3R18 (pink nodes) among which we find Gly, Ala_D, Ser, Glu. Bearing in mind the caveats noted above on the interpretation of the fluxes, it is in principle possible to identify interchangeable strains and their capacities, while also identifying the key metabolites that need to be present in the medium. This opens opportunities to learn how to build new communities or the supplements that a specific strain may need to thrive.

## 3 Discussion

In this study, we introduce `misosoup`, a Python package developed to identify the necessary and sufficient partner species that enable microbial survival in defined environments – referred to as minimal supplying communities (MSCs) and complete minimal supplying communities (CMSCs, see Fig. 1).

`misosoup` builds upon methods used in the SMETANA package [26], which was designed to analyze metabolic interaction potentials between species, summarized as part of the SMETANA score, which aggregates communities and metabolic exchanges. In contrast, `misosoup` focuses on identifying minimal communities that support the growth of all species in the community. This difference in focus is reflected in the outputs of the two packages, and poses additional challenges for `misosoup` in terms of computational complexity. As the communities `misosoup` identifies are used for further analysis, the communities need to be verified to ensure that they do not rely on numerical instability or other artifacts that may arise during the optimization process. This ensures that the communities identified by `misosoup` are biologically meaningful and will be reported consistently. To highlight the complementarity of these packages, we simulated a few examples using both tools (see Suppl. Materials).

Several other relevant packages and algorithms exist, each designed for specific purposes. However, `misosoup` is distinct in several key ways. First, the minimal communities identified by `misosoup` are defined by their ability to support species’ coexistence, rather than being optimized for a purpose such as the synthesis of specific product metabolites [42, 16, 20, 18, 19]. Second, `misosoup` specifically searches for the necessary and sufficient community members required for species’ coexistence, rather than focusing on minimizing the number of species in a community [19, 43] – two different concepts often referred to as “minimal communities”. Third, `misosoup` employs constraint-based methods, unlike graph-theoretic approaches [19, 43, 20, 42], which do not require biomass reactions or precise stoichiometry. This constraint-based approach enhances the accuracy of its predictions.

We explored the potential of `misosoup` to investigate potential applications in biotechnology by first focusing on biogas-producing consortia, finding all MSCs that could grow on ethanol, including those that produce CH4. Notably, we found a previously-reported 3-species consortium, and we illustrated how finding CMSCs allowed us to dig into the modular structure of this consortium. As a limitation, `misosoup` is not designed to search for large communities or to analyze specific communities for (e.g.) optimizing the production of a specific compound, for which other tools are available [16, 35]. Instead, `misosoup` is intended to search MSCs for large sets of species and media, facilitating the exploration of potential consortia and providing guidance on potential assembly rules (e.g. starting from CMSCs and incorporating other strains on top of them).

In another example, “we reproduced results from coculture experiments in which pervasive niche expansion was observed [30]. We found accuracies well above 0.7 for half of the pairs explored, and predicted new ones. There is growing interest in this strategy as shown in another study, where the authors identified 125 instances in which a particular carbon source did not support strain growth in monoculture, but only 21 cases where co-culture also failed to support growth [40]. Thus, the occurrence of no growth in co-culture dropped to 17%, highlighting the potential of this strategy for culturomics.

We also explored the potential of `misosoup` to investigate ecological questions focusing on niche expansion on a larger set of marine strains (60) isolated from experiments of ecological succession of natural communities assembling under controlled conditions [44, 37]. Interestingly, our results are consistent with the ecological strategies identified for these strains, with those found in earlier stages (interpreted in in-vitro experiments with the same strains as degraders and exploiters [45]) being more often suppliers of strains with smaller fundamental niches, which are late colonizers in the experiments. The relationship between facilitation and niche expansion has been previously observed in large-scale studies showing it may be responsible for bacterial cosmopolitanism [46] and community-level coexistence [47]. Although we did not explicitly explore this question, we acknowledge there is growing interest in the evolutionary consequences of the prevalence of facilitation summarized under the Black Queen Hypothesis [48], boosting efforts towards the search of auxotrophies [1, 49, 47] and their role in stability [50].

The importance of facilitation seems to contrast with classical niche theory, which primarily focused on negative biotic interactions, such as predation and competition for space and resources, which constrain an organism’s fundamental niche [51, 52]. In the last two decades, however, there was a shift in focus in macroscopic ecology from the study of competitive exclusion conditions to coexistence conditions [53, 54], highlighting the positive role of mutualistic interactions in promoting stability [55, 56], by reducing the effective competition that species experience [57, 58].

In the microscopic world, there is also growing evidence pointing towards the importance of syntrophy in structuring communities [59]. As a consequence, a large body of theoretical research investigates the role of syntrophy in microbial community stability [60, 61], although with fewer examples showing community-level interaction structures that facilitate coexistence [62, 32, 63]. We illustrated how `misosoup` facilitates the large-scale analysis of communities finding that most syntrophic interactions were bidirectional (cross-feeding).

One reason limiting research in this direction beyond toy models is the computational burden associated with searching and analyzing communities built with metabolic models. In contrast, our approach can, in principle, uncover communities of any size. This capability is due to `misosoup`’s ability to identify minimal communities without exhaustively testing all possible community combinations. For example, with a dataset of 60 strains (as used in this study), there are 1770 possible two-species communities, 34220 three-species communities, and so on, making exhaustive testing impractical if not impossible.

We showed that `misosoup` may help to explore broad theoretical questions such as biodiversity organization, also with potential biotechnological impact. For instance, if there is no need to interact with multiple partners to thrive in a given medium, questions such as how large multispecies communities are sustained remain open. This question connects with how common functional redundancy is in microbial communities, a hot topic in the field [64]. Thanks to the accessible output provided by `misosoup`, it is straightforward to identify functional groups with existing tools (e.g. [41]) by searching for strains consuming and secreting similar metabolites. Identifying strains that may be interchangeable under certain conditions may be invaluable to design new communities and to understand long-standing questions such as how biodiversity is organized from functional groups or how they may combine to build cohesive consortia [31].

Our work has limitations. First, we showed that `misosoup`’s predictive power depends on the quality of the metabolic models. We started with a proof-of-concept example with curated models, then considered semi-curated models for coculture experiments, followed by custom models in the biogas example, and ending with draft metabolic models for the marine communities. Despite `misosoup`’s ability to handle different models, the lower the quality, the less reliable the predictions are and the conclusions drawn from them. Therefore, for broad ecological questions such as the extensive niche expansion observed, it remains to be seen whether it is specific to these species or if it represents a general property of organisms. Although our results with more reliable models also point towards the same conclusion, future studies could replicate our analysis with different sets of species or with random viable organisms that emulate genome-scale metabolic models but contain a random complement of biochemical reactions drawn from a “universe” of such reactions [65].

Second, our insights on the mechanism driving niche expansion come from analyzing a community flux distribution compatible with maximal community growth. However, other objectives (e.g. [66, 67]) might better represent species growth in communities. Exploring these alternatives is straightforward within our framework. `misosoup` can identify the minimal communities (which are not affected by this optimization criterion), and other packages can then be used to find flux distributions aligned with other objectives of interest. In Suppl. Materials we provide an example on how to export `misosoup` results.

Third, despite having the capacity to find communities of any size, the vast majority of communities we found had only 2 members. This could be the consequence of several technical factors specific to our analysis, such as the use of simple media composition, the use of draft metabolic models that could be over-annotated [68], or the number of available strains. Therefore, it remains to be investigated how our results are affected by the total number of species included in the analysis, or the complexity of the medium. As the number of species increases, so does the potential for niche expansion, as the likelihood of including a species with the right set of reactions to complement another one increases. On the other hand, more recalcitrant media may lead to larger minimal communities because more trophic levels may be needed, as described in 45]. The interplay between both, and how large communities are built from minimal ones, remains an open question.

In summary, we developed a package to identify minimal communities, allowing us to explore how communities of any size (not just species pairs) can expand a species’ niche, enabling different potential applications. We hope our tool may contribute to investigating broad ecological questions built around species coexistence, and the development of biotechnological research.

## 4 Methods

### 4.1 Metabolic models

In the following, we briefly describe the models used, with details regarding the generation of mutants and gapfilling procedures available in Suppl. Materials.

First, we tested `misosoup` by performing three analysis with two two-strain and one three-strain consortia, which were experimentally identified to make up minimal communities 33, 21]. For the first analysis with two mutant *E. coli* strains auxotrophic for leucine and lysine 33] we used *E. coli* ‘s metabolic model *i*JO1366 69] which we downloaded from the BIGG database 70]. For the other two analyses, we used the metabolic models of *E. coli, S. enterica* and *M. extorquens* AM1 used in 21] that were provided to us by the authors. For consistency between models, we modified the identifiers of metabolites and exchange reactions (see Suppl. Materials).

For predictions of coculture experiments we considered for simulation three out of five strains assayed in the experiments we attempted to reproduce ([30]), because their metabolic models were available in the AG0RA2 database [34]: *Bacillus subtilis* 3610, *Escherichia coli* B225113, and *Pseudomonas fluorescens* SBW25. We introduced in each model a metabolite id and exchange reaction for each of the 31 carbon sources experimentally assayed, and gapfilled the models to grow on MMAB supplemented with acgam (N-Acetyl-D-glucosamine), because this was the medium they used experimentally to test that all strains were able to grow in monoculture. Further gapfilling was needed for specific carbon sources depending on experimental criteria to determine that growth was observed, detailed in the Suppl. Materials. For consistency, in Main Text we chose the same criteria used by the authors in [30].

To simulate biogas-producing communities the genome-scale metabolic models (GEMs) were directly retrieved from the RedCom publication ([35]). All the models follow the same custom namespace and were not modified.

Finally, we used `CarveMe` [39] to reconstruct the metabolic models for 60 marine species for which whole genome data was available with standard parameters. We simulated marine broth conditions. The detailed composition of the media and the 22 carbon sources selected are available in the Suppl. Materials.

### 4.2 Flux Balance Analysis (FBA)

Flux balance analysis (FBA) is a method to predict metabolic flux – the rate at which chemical reactions convert substrates into products – through every reaction in a metabolic model [10, 9]. It is based on two central assumptions: firstly, that cells optimize some metabolic property such as the growth rate (equal to the flux *r*_*g*_ of an artificial biomass reaction); and secondly, that cells are in metabolic steady-state.

To compute the value of that objective metabolic output, FBA relies on the stoichiometry of every reaction involved in the process. This is mathematically represented as the stoichiometric matrix *S* of size *m* × *v*, where *m* denotes the number of metabolites, and *v* denotes the number of reactions in the model. Each element *S*_*ij*_ corresponds to the stoichiometric coefficient with which metabolite *i* participates in reaction *j*. Additional constraints can be added setting lower (*l*_*j*_) and upper (*u*_*j*_) bounds on the reaction flux (*r*_*j*_). The optimization problem that FBA solves can be written as a linear programming problem:

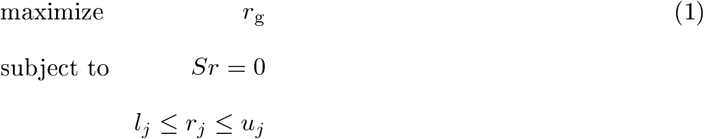

We performed FBA on single strains with `COBRApy` [71].

**Table 1:**
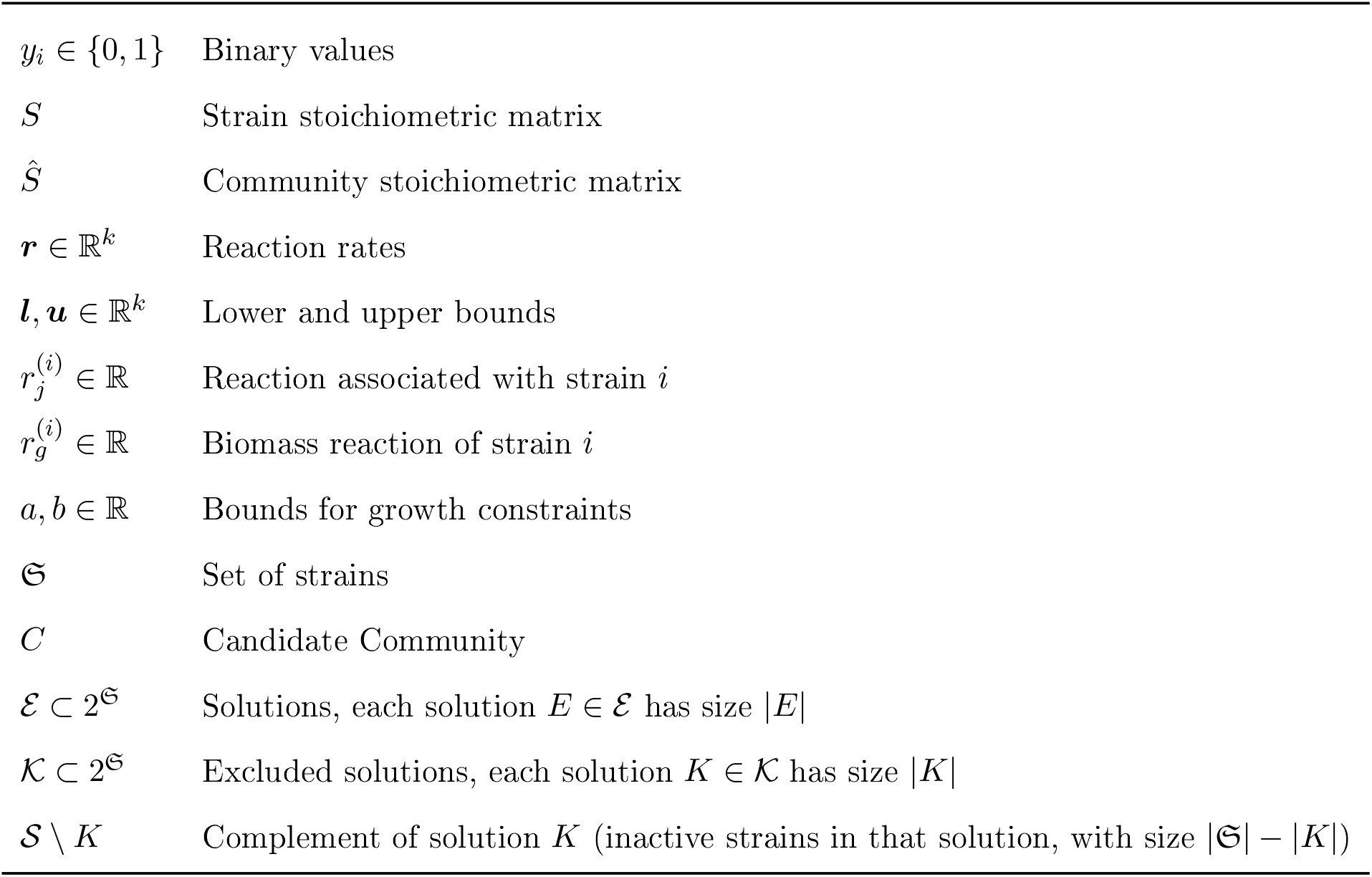
Definitions for the algorithmic MILP / LP problems.

### 4.3 Minimal Community Search

We find minimal communities by searching a larger community metabolic model for minimal sets of strains that can sustain growth. The community metabolic model is constructed by combining all individual metabolic models of the organisms into a single metabolic model with a joint stoichiometric matrix *Ŝ* as in [72]. This formulation allows each organism to exchange metabolites with the environment and with other community members through a shared interface.

Participation of an organism in the community is modeled using a binary variable (*y*_*i*_) as in SMETANA [26]. When a strain is active, the binary variable is set to 1. The strain has to fulfill minimal growth requirements 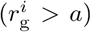 and it can exchange metabolites with the environment and other strains. If the binary variable is set to 0, the strain is inactive and cannot exchange metabolites. Note that a key difference with the work of [26] lies in how the active/inactive state of a strain is modeled. In [26], the sum of secretions is set to zero when a strain is inactive while in `misosoup` the flux through every reaction is set to zero.

Minimal communities are those that minimize the sum of binary variables. We find these by solving the associated MILP problem, similar to SMETANA, by solving the MILP problem specified in Eqs. 2. This problem is constrained by the stoichiometric matrix of the community (*Ŝ*) and lower and upper bounds for each reaction (*l*_*j*_, *u*_*j*_), and bounds on the biomass reaction (*a* and *b*). When a strain is inactive, all its reaction bounds are set to zero, forcing all fluxes to zero.

In addition, two types of constraints are added to progressively restrict the search space. For each accepted community *E* ∈ ℰ, an integer-cut constraint prevents the identical set of active strains from appearing in subsequent solutions. For each discarded solution *K* ∈ 𝒦, we define its complement as 𝒮 \ *K* = {*i* ∈ 𝒮 | *y*_*i*_ = 0}, i.e. the set of strains that were inactive in that solution. A constraint then forces at least one strain in 𝒮 \ *K* to be activated, ensuring that any subsequent candidate community contains at least one strain absent from *K*. These constraints correspond to the last two lines of Eq. 2.

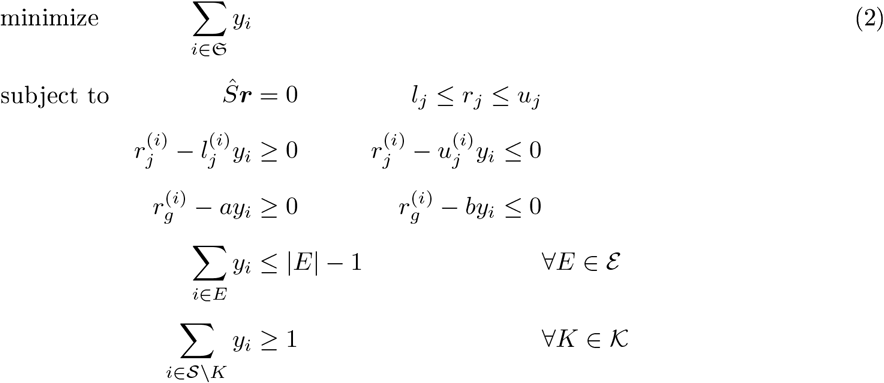

### 4.4 Minimal Community Validation

Solutions to the MILP problem represent potential minimal communities, however, some solutions may be affected by numerical instability inherent to complex problems such as the one we are solving (caused by bigM-constraints for binary variables [73]). In order to verify that the found communities indeed satisfy all constraints imposed (and are therefore feasible solutions), we perform a second optimization with the LP specified in Eqs. 3. For this, we build a community model including only those strains found to compose a potential minimal community 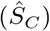 and we maximize the sum of biomass production 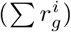 subject to steady-state (*S*_*C*_*r* = 0) and constraints on strain s reaction flux (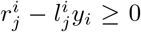 and 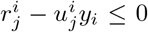) while forcing the strains from the potential community to be in the active state (*r*_*g*_ *> a*). If the optimization is successful, we report the community as a minimal community and add it to ℰ. If the optimization is infeasible, we add the potential community to 𝒦 excluding it from further searches.

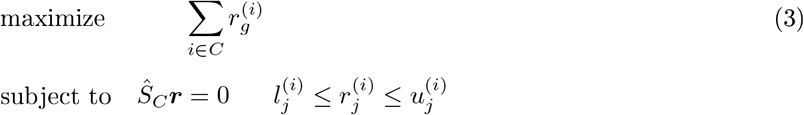

### 4.5 Niche expansion of marine strains

For each marine strain we define its fundamental niche (*F*) as the fraction of environments in which it can thrive in isolation, and its realized niche (*F* ^∗^) as the fraction of environments in which it can thrive in isolation or in a community. The niche expansion (Δ*F*) is defined as the difference between its fundamental niche and its realized niche: Δ*F* = *F* ^∗^ − *F* .

### 4.6 Strains ecological features, phylogeny, and determination of functional groups

We considered the ecological strategies and phylogeny of marine strains inferred from previous work; we provide a brief description here for completeness. Full details can be found in 38]. Ecological strategies of strains were inferred by analyzing the dynamical behavior of Exact Sequence Variants (ESVs) in assembly experiments of marine communities on synthetic particles described in the main text. These experiments had four differentiated compositional stages describing a community level ecological succession. ESV strategies were defined by associating a significant preference to these stages by performing a zero inflated negative binomial regression between each ESV abundance and the factor determining the stages. Strains’ strategies were considered those found for ESVs having a perfect match to them in a 16S rRNA gene alignment.

The phylogeny was also built in previous work [38] using T Tk [7] considering 1 marker genes and formatted in iTOL [75]. Phylogenetic distances were computed from the phylogeny obtained with the function `cophenetic.phylo` of the R package `ape` [76].

Finally, to determine functional groups in Fig. 6 we aggregated all solutions into a single network and analysed it with `functionink` [41]. This algorithm quantifies an all against all similarity between the nodes of a network, where more similar nodes are those connected with approximately the same neighbors with the same type of link (for instance, two species in the same functional group should consume and secrete similar metabolites). To determine functional groups, nodes are clustered according to their similarity, and we considered the maximum of the total partition density as an automatic stopping criterion to determine the optimal partition (see [41] for details).

## Acknowledgements

We thank the group of William Harcombe and in particular to Jeremy Chac ón for providing the metabolic models of mutant *E. coli S. enterica* and *M. extorquens* strains. We also thank members of the Simons Collaboration PRiME, and David San Le ón for useful discussions.

## Data and Code Availability

`misosoup` is freely available at the U L: https://github.com/sirno/misosoup/ and the last release permanently stored in Zenodo with DOI: ‘10.5281/zenodo.15791905’. The 60 genomes of marine isolates are available in NCBI with the BioProject identifiers `PRJNA414740` and `PRJNA478695`. The reconstructed models (in sbml format) and additional data to reproduce results can be found at the U L: https://github.com/sirno/niche_expansion_data stored in Zenodo with DOI: ‘10.5281/zenodo.16744429’.

## 5 Supplementary Materials

### Sl Supplementary Methods

#### S1.1 Modification of identifiers in metabolic models

We modified the identifiers of metabolites and exchange reactions only to ensure consistency between models, and they are not required for the use of `misosoup`.

In models used in the first analysis of 2- and 3-species consortia, we replaced every compartment indicator `[e]`, `[p]` and `[c]`; or `(e)`, `(p)` and `(c)` by `_e`, `_p` and `_c` respectively. Further, in S. enterica’s model we changed the identifiers of the following external metabolites (`<old>/<new>`): `cam/cm`, `galctr_ _D/galct_ _D`, `glcn_ _D/glcn`, `metox_ _R/metsox_ _R_ _L`, `metox/metsox_ _S_ _L`, and `orn_ _L/orn`. Similarly, we changed the identifiers of two external metabolites in *M. extorquens’* model: `glcn_ _D/glcn` and `mea/mma`. Additionally, we changed the biomass reaction identifier from `BI02b_Mex` to `Me_biomass`. Last, we changed the identifiers of every exchange reaction in all three models so that they were of the form `EX_<identifier>`. For example, the external reaction corresponding to glucose (with metabolite id `glc_ _D_e`) is of the form `EX_glc_ _D_e`.

#### S1.2 Construction of mutants

For *E. coli* ‘s metabolic model *i*JO1366 [69] used in the first example, we modified it to build the respective mutant strains: for mutant lysA, we constrained flux through the associated reaction (DAPDC) to zero; for mutant leuA, we constrained flux through the associated reaction (IPPS) to zero.

In the example predicting coculture experiments, we created models for auxotrophic strains from WT gapfilled models for leucine (L), arginine (R), tryptophan (W) and histidine (H) amino-acids. For this, we blocked in the models every reaction that biosynthesizes the specific amino-acid. We also blocked catabolic pathways using these amino-acids. This means that we set lower and upper bounds to zero for every reaction in which the metabolite was either produced or consumed. We took this rather conservative approach because, experimentally, they used different ways to generate auxotrophies which may not be in reproducible in the models. For instance, Oña’s *et al* . [30] knocked out specific genes (argH, leuB, trpB or trpC and hisD) and found the desired phenotype, while in the models there were alternative enzymes. Indeed, *E*.*coli* and *P*.*fluorescens* WT models were able to grow with the amino-acid as a sole carbon source, a behavior that was not observed in the experiments.

**Table S1:** Metabolite composition of the MMAB base media.

| ID | Name | Formula/Ion |
| --- | --- | --- |
| ca2 | Calcium | $\text{Ca}^{2+}$ |
| cl | Chloride | $\text{Cl}^-$ |
| cobalt2 | Cobalt | $\text{Co}^{2+}$ |
| cu2 | Copper | $\text{Cu}^{2+}$ |
| fe2 | Iron (II) | $\text{Fe}^{2+}$ |
| k | Potassium | $\text{K}^+$ |
| mg2 | Magnesium | $\text{Mg}^{2+}$ |
| mn2 | Manganese | $\text{Mn}^{2+}$ |
| na1 | Sodium | $\text{Na}^+$ |
| nh4 | Ammonium | $\text{NH}_4^+$ |
| o2 | Oxygen | $\text{O}_2$ |
| pi | Phosphate | $\text{PO}_4^{3-}$ |
| so4 | Sulfate | $\text{SO}_4^{2-}$ |
| zn2 | Zinc | $\text{Zn}^{2+}$ |
| h2o | Water | $\text{H}_2\text{O}$ |
| h | Protons | $\text{H}^+$ |

#### S1.3 Detailed composition of media

##### Coculture experiments

In coculture experiments, composition of MMAB base media is displayed in Suppl. Table S1, and it was available in unlimited amounts. In addition, each of the 31 C sources considered in biolog plates are shown in Suppl. Table S2.

##### Marine medium

We simulated a marine environment assuming that ammonium, biotin, calcium, chloride, cobalamin, cobalt, copper, folate, iron, magnesium, manganese, molybdate, nickel, nicotinic acid, oxygen, p-aminobenzoic acid, pantothenic acid, phosphate, potassium, protons, pyridoxal, riboflavin, selenate, selenite, sodium, sulfate, thiamine, tungstate, water, and zinc were available in non-limiting amounts.

For each marine environment a single carbon source was available at a time. The 22 carbon sources used in the analysis were selected through a model-based screening approach. First, Flux Balance Analysis (FBA) simulations were performed in marine medium using each potential carbon source individually. Potential carbon sources were defined as carbon-containing metabolites associated with an exchange reaction in the model (410 metabolites in total), and simulations were conducted across all strains included in the study. For each substrate, we calculated the fraction of strains predicted to grow. From this set, we selected 22 carbon sources listed in S3 spanning a broad range of growth prevalence across strains (Fig. 5C in Main Text, black bars), while maintaining a balanced representation of major compound classes, including carboxylic acids, amino acids, and carbohydrates.

**Table S2:**
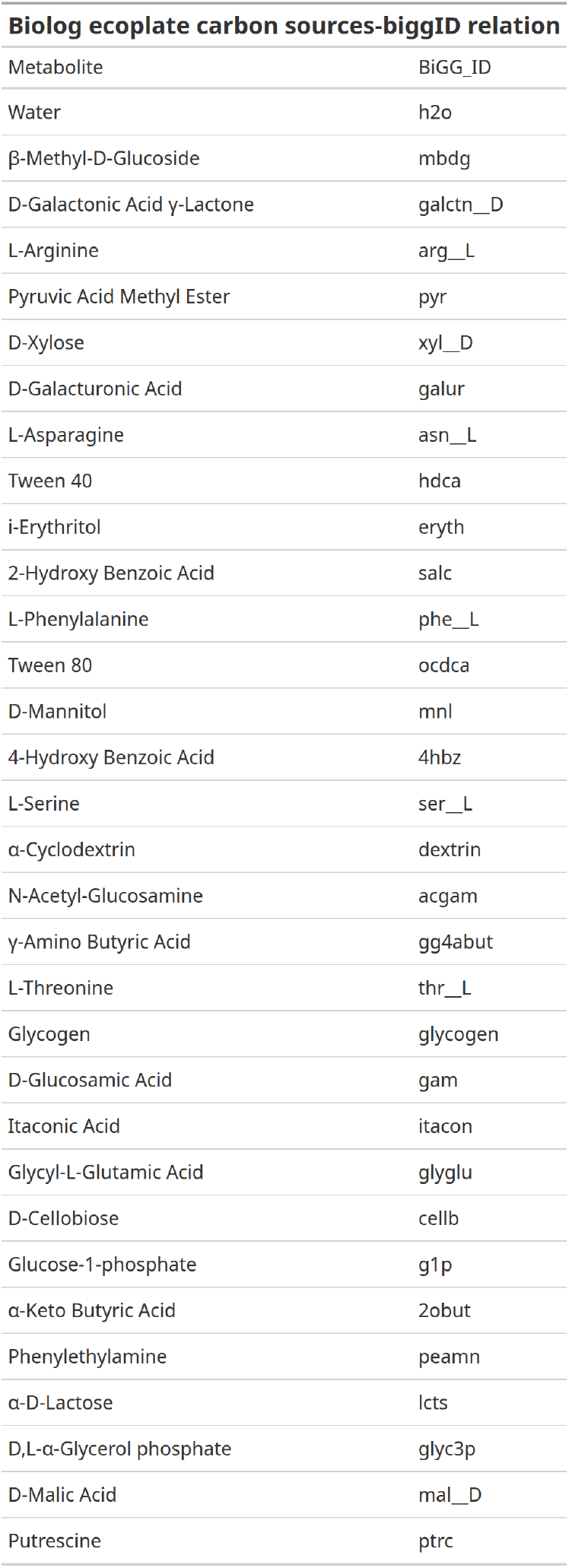
Carbon sources of the Biolog Ecoplates and its BIGG ids. In the metabolites which does not have a direct ID, proxy metabolites where selected.

**Table S3:** Carbon sources used in marine environments.

| ID | Name | Formula |
| --- | --- | --- |
| pep | Phosphoenolpyruvate | $\text{C}_3\text{H}_2\text{O}_6\text{P}^{2-}$ |
| icit | Isocitrate | $\text{C}_6\text{H}_5\text{O}_7^{3-}$ |
| glu__D | D-Glutamate | $\text{C}_5\text{H}_8\text{NO}_4^-$ |
| 3pg | 3-Phospho-D-glycerate | $\text{C}_3\text{H}_5\text{O}_7\text{P}^{2-}$ |
| r5p | Ribose 5-phosphate | $\text{C}_5\text{H}_9\text{O}_8\text{P}^{2-}$ |
| met__L | L-Methionine | $\text{C}_5\text{H}_{11}\text{NO}_2\text{S}$ |
| akg | 2-Oxoglutarate | $\text{C}_5\text{H}_4\text{O}_5^{2-}$ |
| oaa | Oxaloacetate | $\text{C}_4\text{H}_2\text{O}_5^{2-}$ |
| f6p | D-Fructose 6-phosphate | $\text{C}_6\text{H}_{11}\text{O}_9\text{P}^{2-}$ |
| phe__L | L-Phenylalanine | $\text{C}_9\text{H}_{11}\text{NO}_2$ |
| trp__L | L-Tryptophan | $\text{C}_{11}\text{H}_{12}\text{N}_2\text{O}_2$ |
| rmn | L-Rhamnose | $\text{C}_6\text{H}_{12}\text{O}_5$ |
| for | Formate | $\text{CHO}_2^-$ |
| ile__L | L-Isoleucine | $\text{C}_6\text{H}_{13}\text{NO}_2$ |
| rib__D | D-Ribose | $\text{C}_5\text{H}_{10}\text{O}_5$ |
| ser__D | D-Serine | $\text{C}_3\text{H}_7\text{NO}_3$ |
| acgam | N-Acetyl-D-glucosamine | $\text{C}_8\text{H}_{15}\text{NO}_6$ |
| cit | Citrate | $\text{C}_6\text{H}_5\text{O}_7^{3-}$ |
| ac | Acetate | $\text{C}_2\text{H}_3\text{O}_2^-$ |
| pyr | Pyruvate | $\text{C}_3\text{H}_3\text{O}_3^-$ |
| glc__D | D-Glucose | $\text{C}_6\text{H}_{12}\text{O}_6$ |
| pro__L | L-Proline | $\text{C}_5\text{H}_9\text{NO}_2$ |

#### Sl.4 Gapfilling metabolic models for coculture experiments’predictions

In coculture experiments performed by (Oña et al. [30]), the authors explored niche expansion for different strains comparing their growth in monoculture and in coculture. They considered both wild type strains, and auxotrophic strains for four amino-acids (leu, arg, trp and his). They grew these strains in MMAB having one of the 32 carbon sources available in biolog ecoplates, with and without supplementation in the media of the KO amino-acid.

##### Influence of experimental criteria determining strains’ growth

For each culture, they provided measurements of optical density (OD) at different days (samples taken after 0, 3, 5 and 7 days of growth) in triplicate. Importantly, they considered growth if at least one out of three biological replicates had an OD above 0.08. As we will show below, the accuracy of `misosoup` necessarily depends on these choices, so we systematically explored its accuracy as a function of variations in these parameters.

To compare the results with the experiment, we built for each combination of experimental criteria parameters *c* (OD threshold, number of replicates, and time), two binary vectors 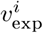 and 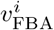 where *I* corresponds to each of the acgam-gapfilled models of the three strains, describing growth (= 1) or not (= 0). By comparing these vectors, we estimate the extent to which the models perform like the experiments, what will affect downstream analysis.

The first test we conducted was to verify that, whenever growth was observed in monoculture experiments, the acgam-gapfilled models were able to reproduce it. Oña *et al*. considered as a criterion to determine growth, that an OD of 0.08 should be observed in at least one replicate. Therefore, we verified whether variations in this criteria influenced predictive performance. We considered different combinations *c* of OD thresholds, number of replicates and growth times to determine performance in acgam-gapfilled models. We found that, at day 0, the models showed high sensitivity for low thresholds when growth in 1 or 2 wells was considered, whereas sensitivity is low when 3 wells was considered, also having low specificity (Fig.S1). This behaviour is expected, because bacteria had no time to grow. Hence, when compared with models’ predictions, it translates into a high number of false positives and a low number of false negatives, hence lowering specificity but increasing sensitivity. In the remaining scenarios, the number of wells is the only factor that increases sensitivity, with no systematic impact on specificity.

**Figure S1:**
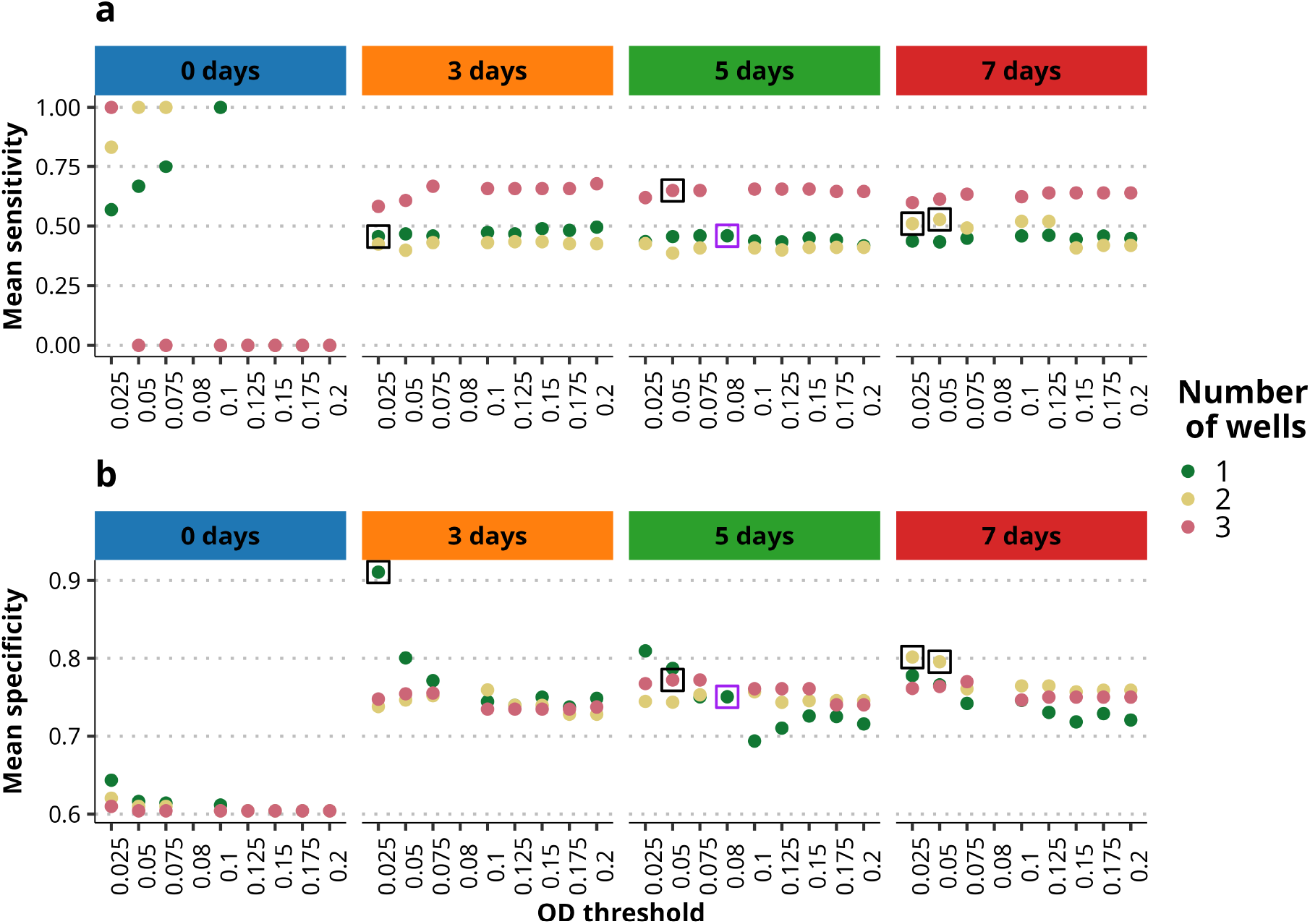
Mean performance of metabolic models for different combinations of experimental parameters. Performance was tested using acgam-gapfilled models. Selected combinations are indicated with black squares, with a purple one for Oña’s *et al* choice. The number of wells in which growth was observed is the factor that has the greatest influence on sensitivity, whilst the impact on specificity depends to a greater extent on the optical density from the third day onwards. The remaining factors do not affect the performance with the exception of time at day 0, where cultures have not achieved high ODs.

##### Influence of gapfilling in WT metabolic models performance

Since criteria to determine whether a species grows determines the media in which models should be gapfilled, we explored the effect of gapfilling depending on the selected combinations of parameters. We selected five *c* combinations, indicated with squares in Fig. S1. We chose one combination in which there was growth in all three wells with an intermediate OD, as replicability was the factor that most notably increases sensitivity. Then, we selected three scenarios with high specificity, because we expect gapfilling to preferentially increase sensitivity. Finally, we selected the scenario chosen by Oña *et al*. (purple square).

**Figure S2:**
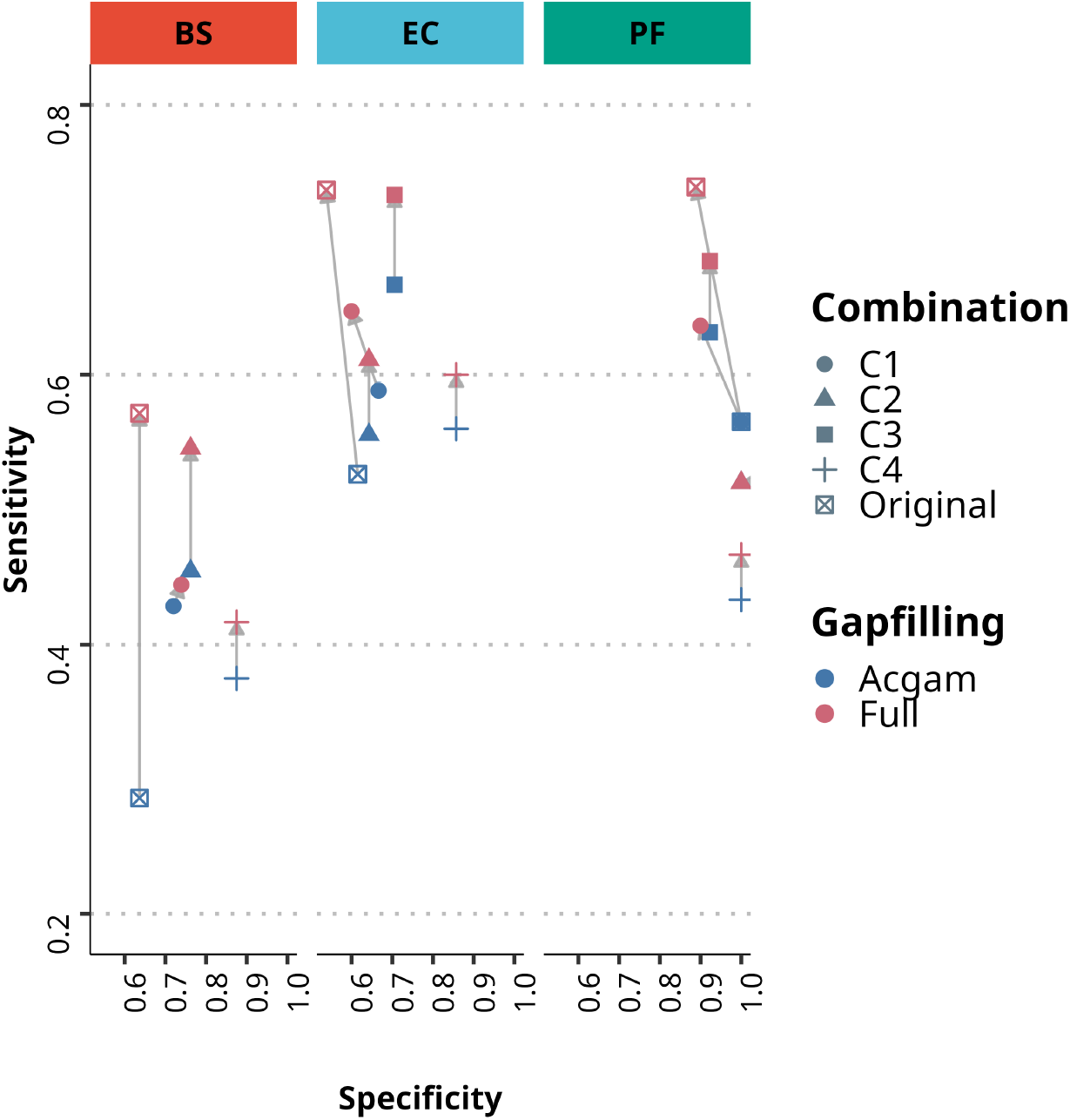
Performance of selected experimental parameter combinations before and after gapfilling WT models. Sensitivity and specificity for each acgam-gapfilled model before and after gapfilling show that gapfilling improved sensitivity, with little effect on specificity. BS had a notably lower initial sensitivity, suggesting the quality of the model was poorer. Despite improvement after gapfilling, it very likely affected its performance in coculture (also see Suppl. Fig. S4).

We gapfilled WT models considering the 5 combinations of experimental parameters selected. For this1 we considered acgam-gapfilled WT models to explore which carbon sources considered in the experiment `misosoup` predicted growth when growth was indeed observed. Growth was determined using FBA with `cobrapy`, using the method `.optimize()`, and considering that the objective function value should be above 0.01. For each carbon source in which there was growth in the experimental data but not in our model, we further gapfilled the model for that specific C source using the gapfilling utility available in `carveme`. However, for 8 out of 32 C sources gapfilling led to thermodynamically infeasible cycles in which models were able to generate ATP without any source of energy in the medium. These C sources were itacon, ocdca, 4hbz, peamn, galctn D, glycogen, and eryth, so no gapfilling was considered in these media

As a final note, some carbon sources had no direct ID in BIGG, so we selected similar metabolites whenever possible (for example, for tween 40 or tween 80 the fatty acids without the ester were used as a proxy). In addition, ciclodextrin was not included since there was no reliable proxy in BiGG. This led to experiments on 31 carbon sources. After gapfilling, models’ sensitivity improved with no effect on specificity, as expected (see Fig.S2). The combination selected by the authors (indicated as ‘original’) was the one showing a higher increase in sensitivity. Note that, in C3, there was no growth for BS. Therefore, since there was no strong evidence of improvement to choose another scenario, for consistency with Oña’s results we continued our analysis using the original combination of parameters.

##### Influence of misosoup’s objective function values in its accuracy

Finally1 since `misosoup` predictions may also depend on the specific community growth threshold1 *g*_c_, we analysed whether it influences `misosoup` accuracy. For this analysis1 we considered the ‘original’ combination of experimental parameters1 although other choices led to qualitatively similar results (see below).

As was the case for individual models, to compare the results with the experiment we built, for each combination *c* of parameters determining whether growth was observed, two binary vectors *v*^exp^ and *v*^mis^, describing observation (= 1) or absence (= 0) of growth in the experiment and in `misosoup` predictions, respectively.

We ran `misosoup` with the MSC algorithm without any community growth threshold in its version 2.3.0. e simulated all against all combinations of the four auxotrophs (H, R, L, W) for the 3 strains in 31 substrates1 leading to 2046 simulations. To focus on `misosoup` performance and neglect errors that can objectively be attributed to the use of automatically curated models, `misosoup` accuracy was computed on cases in which there was an agreement between experiments and prediction for auxotrophic strains grown in monoculture, when the amino-acid missing was supplemented in the medium.

Increasing *g*_c_ led to an almost constant trend in sensitivity and specificity until a threshold around 0.6 where there was a sharp drop in sensitivity and a less pronounced increase in specificity. Since, in light of the accuracy, the increase in specificity doesnot compensate the decrease in sensitivity, we conservatively chose a threshold of 0.5 for downstream analysis.

**Figure S3:**
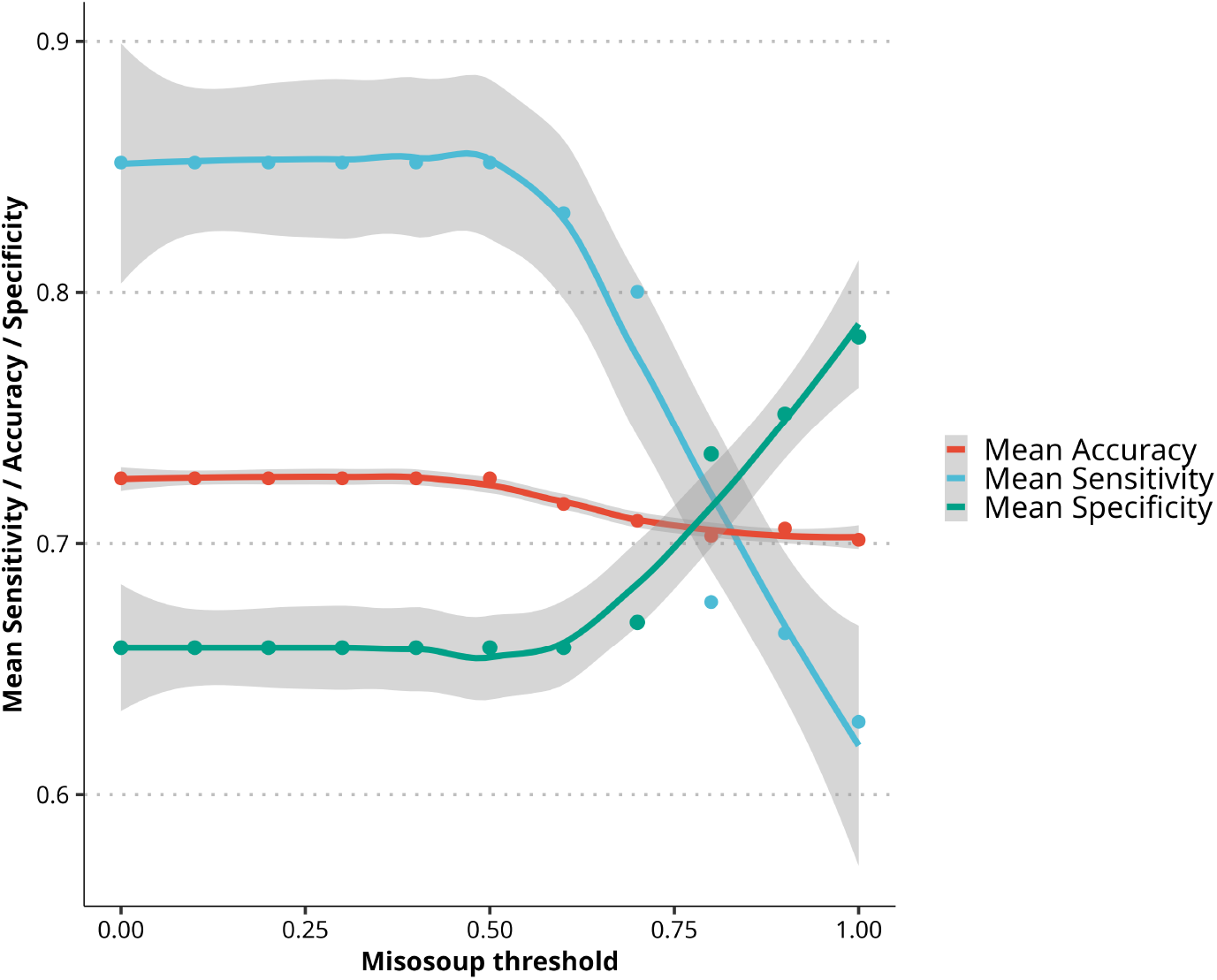
Performance of misosoup for different community growth thresholds. Although the mean accuracy is almost constant across the values of *g*_c_; however, there is a clear trade-off between sensitivity and specificity for values of *g*_c_ above 0.75 with the accuracy remaining constant along the different community growth thresholds.

##### Consistency of gapfilling with Carveme

We tested whether gapfilling in `Carveme` depends on the specific run when several substrates were considered for gapfilling. After running 10 iterations for each model, we found that the reactions added were not identical in each iteration, although the ability of the final model to grow on the selected substrates remained the same. There was a set of core reactions added in all iterations and a set of variable reactions Fig.S4. EC and PF were more consistent across iterations than BSwas. Moreover, in one of the iterations excluded in Fig. S4), gapfilling of BS introduces an infeasible cycle due to the addition of the 0tex transporter, which allowed the model to remove protons from the cell at no cost. These results pointed once again towards a lower quality of this model, which likely explains the higher number of false positives found in coculture experiments when it was included in the prediction.

#### Sl.5 Working with minimal communities outside misosoup

The minimal communities identified by `misosoup`can be further analyzed using additional software packages. This is particularly useful when aiming to explore consumption and secretion profiles of each strain in the community under objectives beyond those implemented in `misosoup`(e.g., maximizing community biomass or minimizing total flux). Here, we provide an example of constructing a community model for a minimal community comprising three species and exporting it as an SBML file for compatibility with other tools.

**Figure S4:**
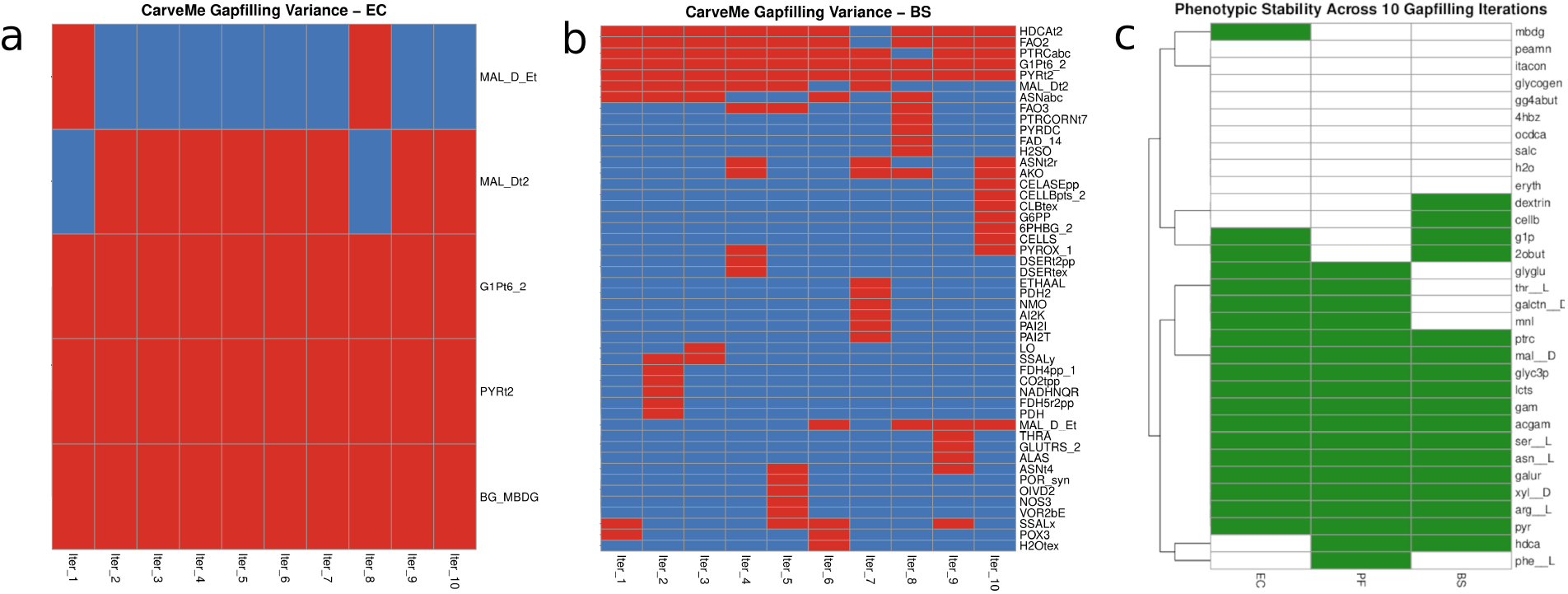
Gapfilling consistency. `Carveme` does not consistently add the same reactions when different runs are performed to gapfill the same media. Reactions added (red cells) in each of the 10 iterations performed for *E. Coli* (A) and *B. Subtilis* (B). BC model required gapfilling in more media, indicating that the model initially had a lower quality. Despite this variability, phenotypic plasticity was consistent (C), with models consistently growing in the same media in all 10 iterations (green cells). EC = *E. Coli* ; PF = *P. fluorescens*; BS = *B. Subtilis*.

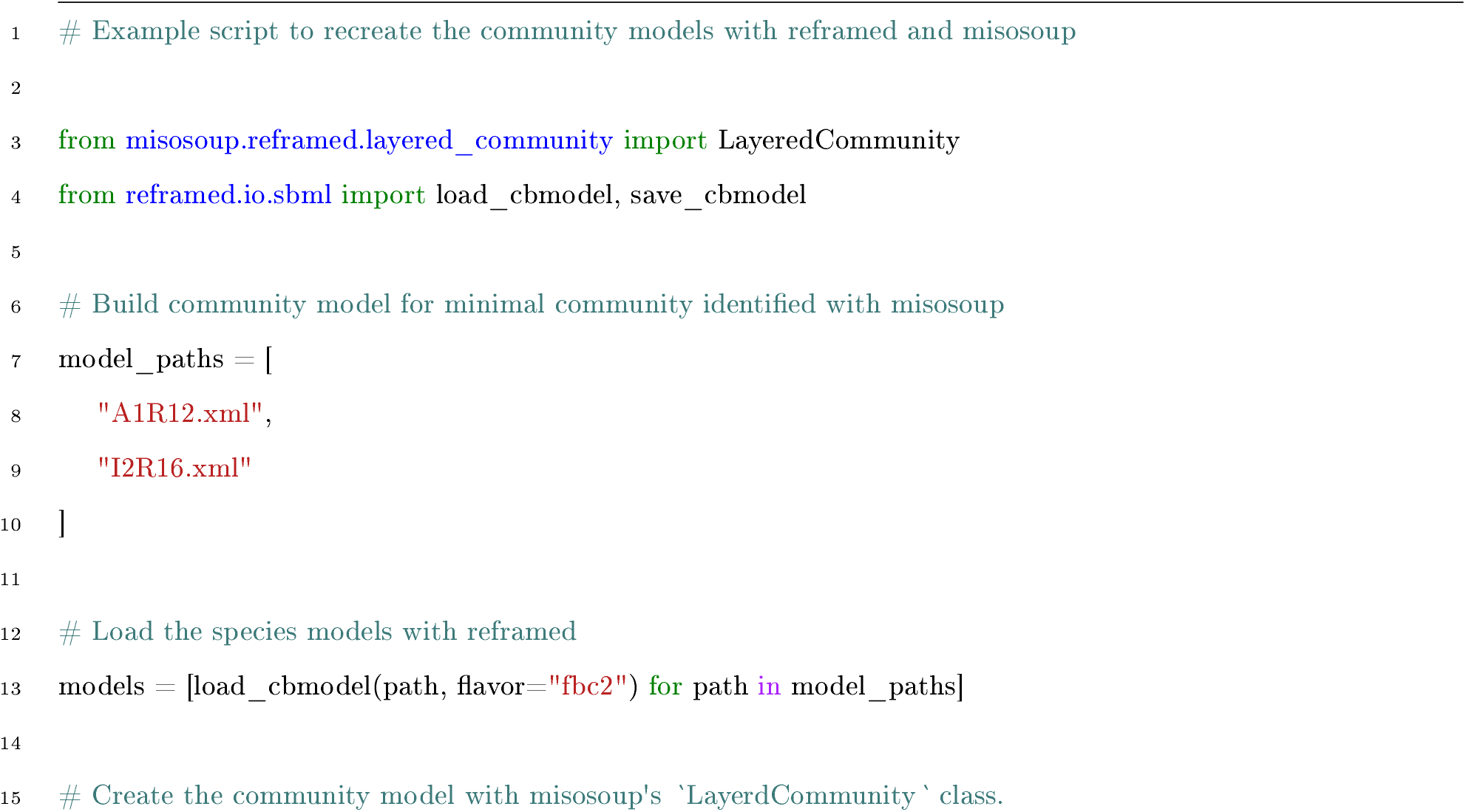
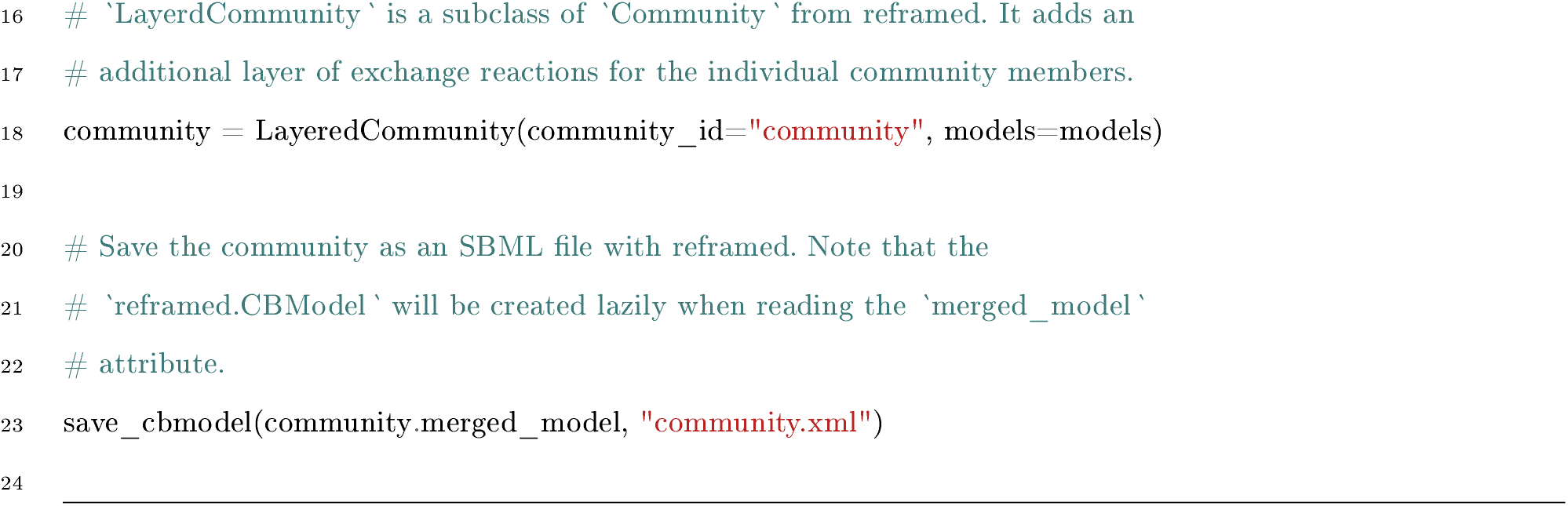

### S2 Supplementary Results

#### S2.1 Supplementary Note. Comparison of packages SMETANA and misosoup

To illustrate the difference between the analysis performed by SMETANA [26] and `misosoup`, we simulated two examples for each method. Results can be found in https://github.com/sirno/niche_expansion_data under the path: `data/examples/comparison_smetana_misosoup`.

For the first example we used strains Arcobacteraceae_A1R12, Alteromonadaceae_I2R16 and Vibrionaceae_I3M07 from the marine strains’ dataset. First, we used `misosoup`to search for minimal supplying communities that would allow growth of Arcobacteraceae_A1R12 (the focal strain) in marine medium (see Method’s section S1.3) with acetate as the sole source of carbon. We found two MSCs, with Alteromonadaceae_I2R16 or Vibrionaceae_I3M07 as supplying strains. `misosoup`’s output returns the minimal supplying communities found, as well as a detailed description of each species consumption/secretion pattern and growth rate, compatible with maximal community growth.

Then, we used the same three species and medium composition and used SMETANA, in both global and detailed modes (see `example1/smetana/run.sh`). The global analysis returns two scores: metabolic interaction potential (mip) and metabolic resource overlap (mro). These quantify the number of nutritional requirements that the community can provide through cross-feeding interactions, and the overlap in the minimal nutritional requirements, respectively. The detailed analysis returns three scores (scs, mus and mps) which are combined into a fourth score named SMETANA that quantifies the cross-feeding plasticity between the community members.

Note that for this first example, we could have inferred the two minimal supplying communities obtained with `misosoup`from the detailed analysis of SMETANA. In `example1/smetana/output_exam ple1_detailed.tsv` we find that both Alteromonadaceae_I2R16 and Vibrionaceae_I3M07 can act as donors of metabolites which are required for growth of Arcobacteraceae_A1R12. However, this can only be easily deduced when no more than two strains participate in the interaction.

In the second example (see `example2/`) we used a set of three other strains (Vibrionaceae_1A01, Psychromonadaceae_B3M02 and Alteromonadaceae_D2M02) and simulated a marine medium (see Method1s section S1.3) with D-glutamate (`glu_ _D`) as carbon source. First, we used `nisosoup`to find a minimal supplying community that would allow growth of Psychromonadaceae_B3M02. We found that only when both Vibrionaceae_lA0l and Alteromonadaceae_D2M02 are present, Psychromonadaceae_B3M02 grows. In contrast, when we used SMETANA for the analysis of this community, both, the global and detailed analysis will not contain any results, because on this medium, none of the three strains can grow in isolation.

#### S2.2 Supplementary Note. Additional results for coculture experiments

**Figure S5:**
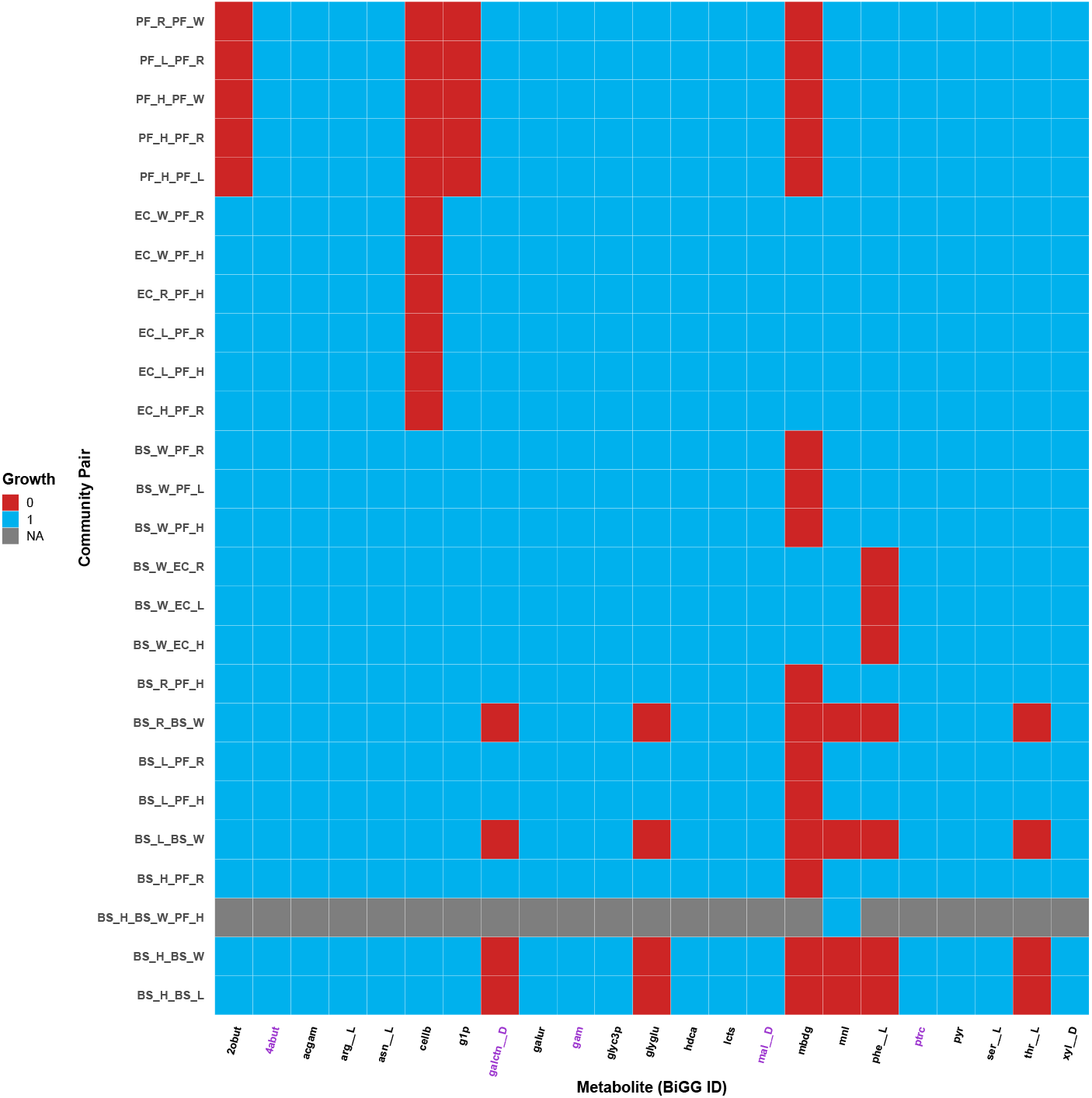
Additional coculture predictions obtained with `misosoup` that were not experimentally verified. Those involving BS should be taken with caution, given the false positives detected in previous analysis. Interestingly, `misosoup` also detects coculture pairs formed by pairs of auxotrophs of the same species.

#### S2.3 Supplementary Note. Additional results for marine strains simulations

##### MSCs show high metabolic and niche complementarity with phylogenetic relatedness at or above the Family level

As we noted in Main Text, we found that the lower the phylogenetic distance, the less similar the reactions are between pairs of species (Suppl. Fig. S6, left), which translates into an also lower niche similarity (Suppl. Fig. S6, right). In addition, we also observed that, if the fundamental niche size is larger than zero, the higher the number of genes or reactions, the higher the fundamental niche size (Suppl. Fig. S7).

**Figure S6:**
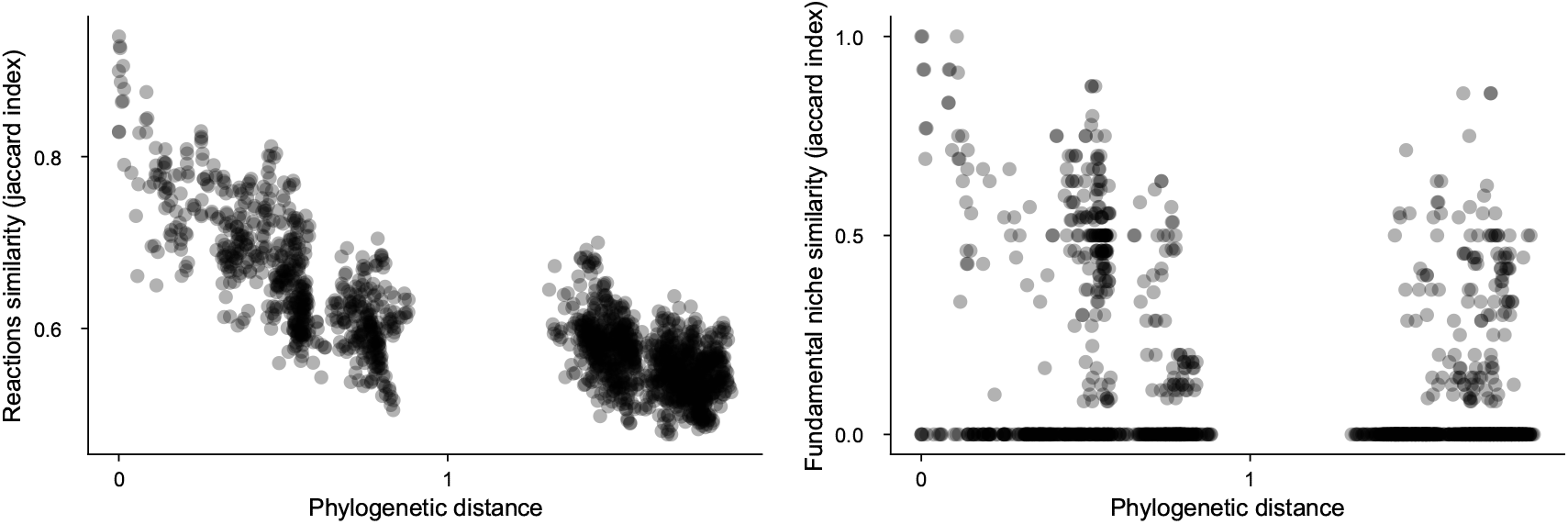
Relationship between phylogenetic distance and reactions similarity (left) and fundamental niche similarity (right). Each point represents a pair of strains. Reaction similarity is calculated as the Jaccard index, defined as the ratio of shared reactions to the total number of unique reactions (union of reactions) .

**Figure S7:**
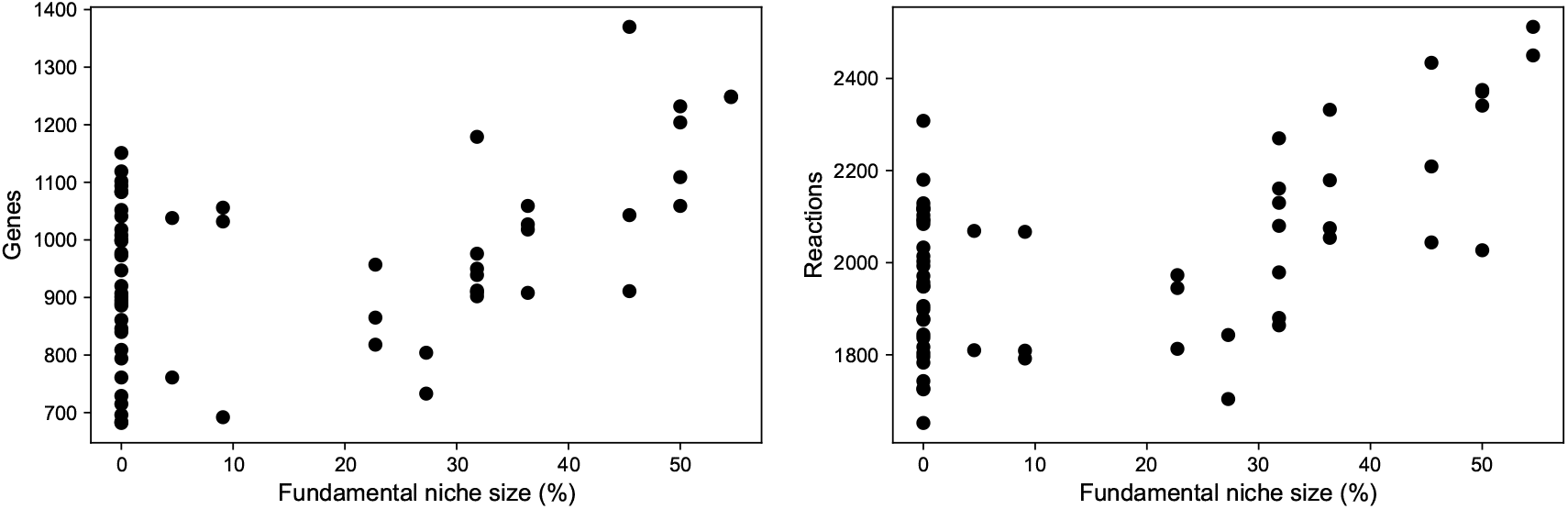
Species’ fundamental niche vs number of genes (left) and reactions (right) in its metabolic model. Niche size is quantified as percentages of environments that a species thrives in.

Digging further into these observations, we investigated the phylogenetic relationships among members of minimal supplying communities (MSCs), initially focusing on 2-strain communities. We found that the strains forming MSCs predicted by `misosoup` are, on average, phylogenetically closer to each other than randomly selected strain pairs (Fig. S8A). Importantly, while MSCs showed increased phylogenetic relatedness compared to random communities, they were not exclusively composed of members from the same family – communities composed of strains from different families are also frequently observed. This general trend of phylogenetic relatedness held true for 3-strain MSCs as well, which were similarly composed of phylogenetically closer strains than randomly formed groups (Suppl. Fig. S10).

In [30], the authors investigated whether the overlap in strains’ fundamental niches could be used to predict the degree of niche expansion. Specifically, for every pair of strains they quantified the union minus the intersection (hereafter referred to as the symmetric difference) of the strains’ fundamental niches. They found that, indeed, the symmetric difference between strains’ fundamental niches could predict the degree of niche expansion, where a larger symmetric difference is indicative of a larger niche expansion.

Building on this observation, we calculated the symmetric difference of the fundamental niches of strains that constitute the predicted minimal supplying communities, and then compared these to the symmetric differences found in randomly selected sets of strains from the same species pool. We hypothesized that strains forming MSCs identified by `misosoup` would exhibit a larger symmetric difference in their fundamental niches.

Our findings confirmed this hypothesis. Consistent with results in [30], we observed that the symmetric difference of fundamental niches for the predicted communities was higher than that of random communities (Fig. S8B). Interestingly, this distinction between MSCs and random communities was primarily driven by a larger union in the fundamental niches (Suppl. Fig. S11) rather than by a smaller intersection.

Based on this observation we decided to investigate whether the differences between random and predicted communities appeared at genetic levels. For each minimal community, we considered the set of genes (or reactions) of each strain *G*_*i*_ and computed the union, intersection and symmetric differences of pairs or triplets of strains. For instance, the symmetric difference between two strains *i* and *j* is defined as the union minus the intersection *G*_*i*_Δ*G*_*j*_ = (*G*_*i*_ ∪ *G*_*j*_)*/*(*G*_*i*_ ∩ *G*_*j*_). We tested the values found by building a random distribution by selecting pairs of strains randomly with replacement. Again, we observed that there was a significant difference between predicted and random communities, with random communities showing a lower symmetric difference in the metabolic reactions S8C.

Overall, we observed that MSCs were made up of pairs of strains from the same or different family with complementary sets of reactions. This complementarity in metabolic reactions is likely to explain the complementary fundamental niches of the strains involved. Importantly, this complementarity of metabolic reactions and fundamental niches is not observed in communities created at random from the same pool of strains.

**Figure S8:**
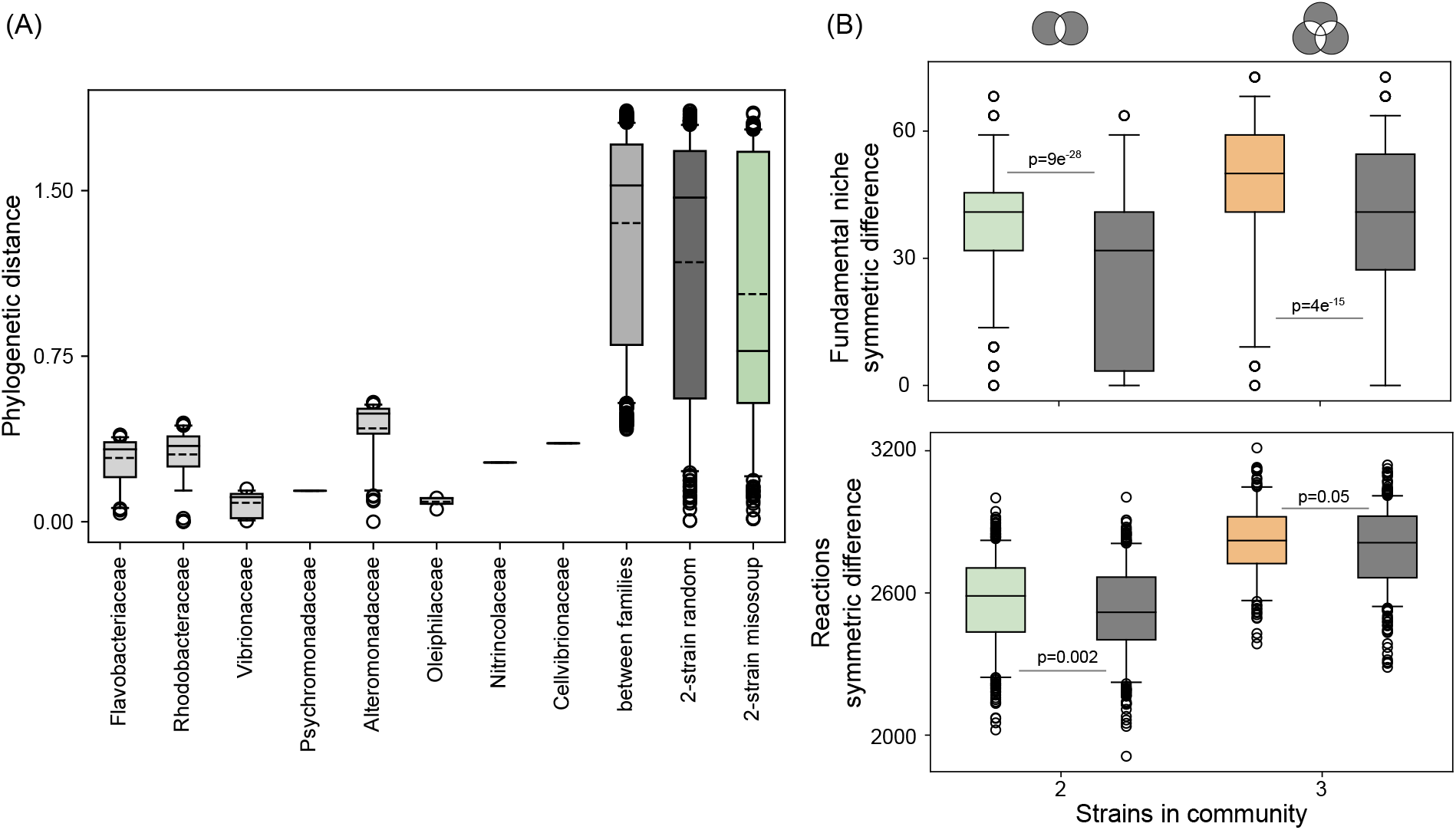
(A) Phylogenetic distances between pairs of strains within the same family (family names indicated on the x-axis), between pairs of strains from different families, between 500 randomly selected strain pairs, and between 2-strain communities predicted by `misosoup`. Symmetric difference of fundamental niches (B) and reactions in the metabolic models (C) for MSCs and 500 communities of random composition. Results for MSCs with two or three strains are shown in green and orange respectively. Results for random communities are shown in gray. Boxes extend from the first quartile to the third quartile of the data, with a line at the median.

##### Supplementary figures from marine strains simulations

**Figure S9:**
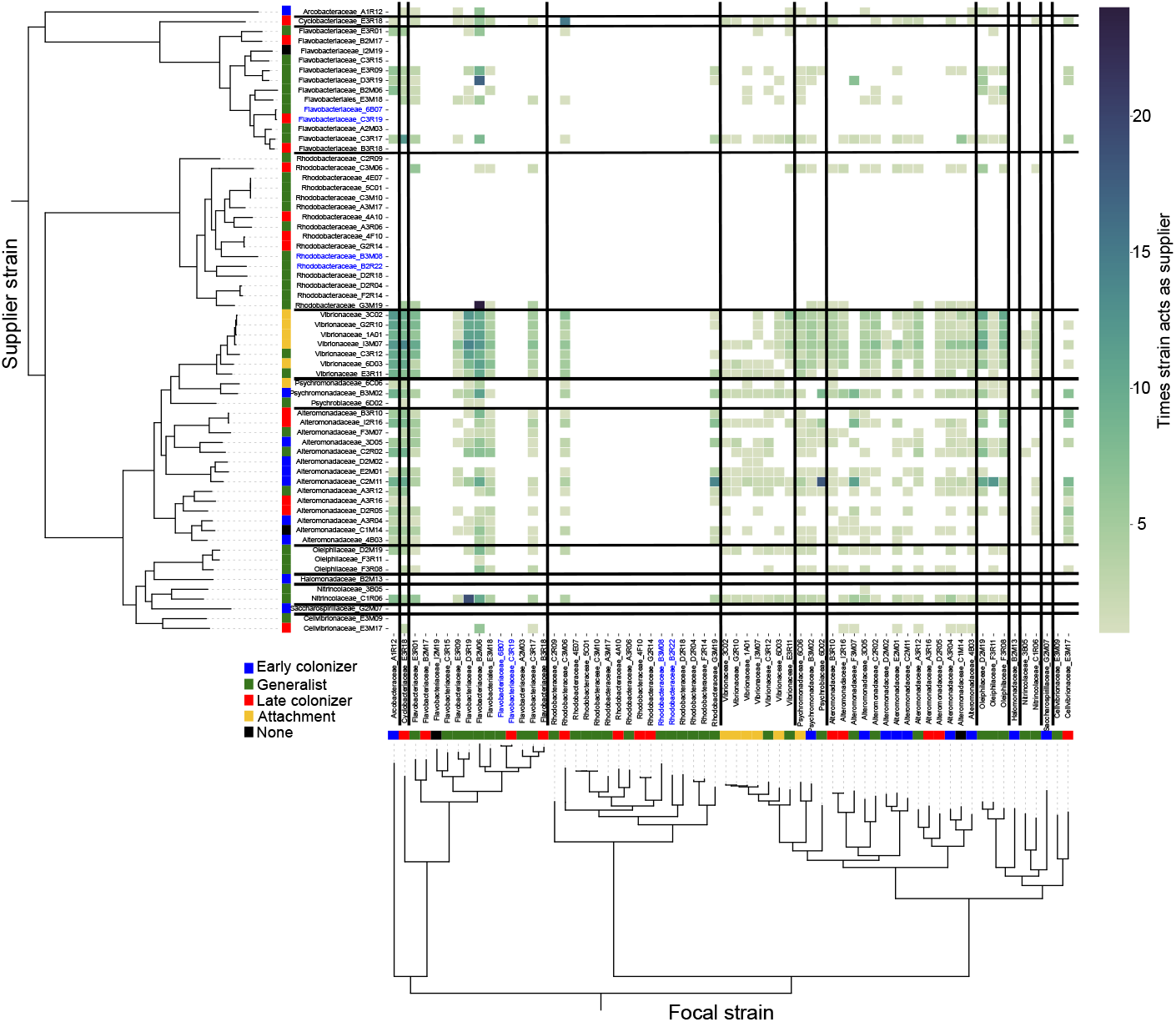
Heatmap depicting the times a strain acts as a supplier (y-axis) to a focal strain (x-axis). The phylogenetic tree and ecological strategies of the strains, as inferred in Pascual-Garcia (2022), are shown for both focal and supplier strains. Black lines indicate boundaries between strains from different families.

**Figure S10:**
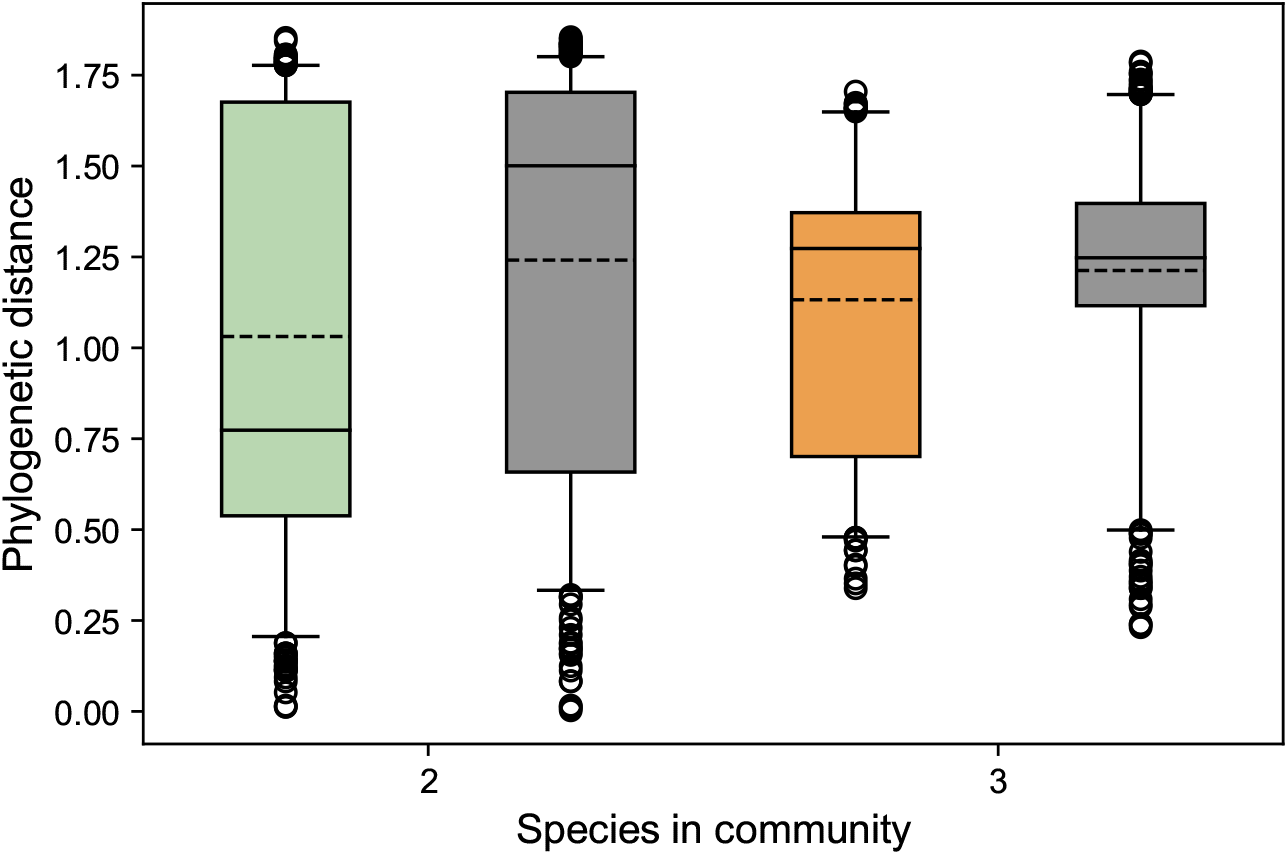
Phylogenetic distance for MSCs and 500 communities of random composition. Results for minimal supplying communities with two or three species are shown in green and orange respectively. Results for random communities are shown in gray. For communities made up of three species, the phylogenetic distance shows the average distance among the three possible pairs of species . Boxes extend from the first quartile to the third quartile of the data, with a communities line at the median and a dashed line at the mean. Phylogenetic distance in MSCs is significantly smaller than that of communities built at random from the same species pool.

**Figure S11:**
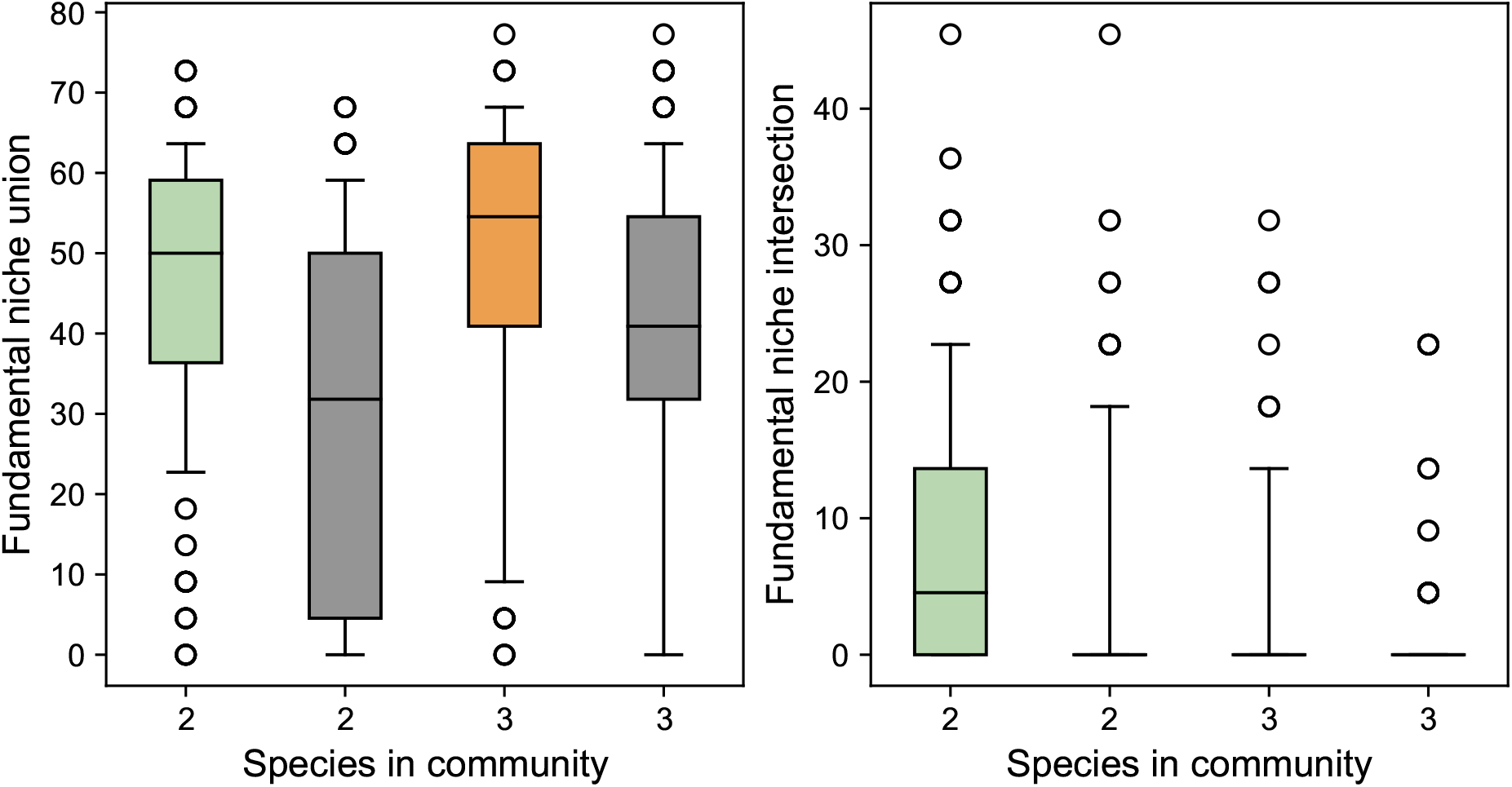
Fundamental niche union (left) and intersection (right) for minimal supplying communities and 500 communities of random composition. Results for MSCs with two or three species are shown in green and orange respectively. Results for random communities are shown in gray. Boxes extend from the first quartile to the third quartile of the data, with a line at the median.

